# MetaIBS - large-scale amplicon-based meta analysis of irritable bowel syndrome

**DOI:** 10.1101/2024.01.22.575775

**Authors:** Salomé Carcy, Johannes Ostner, Viet Tran, Michael Menden, Christian L. Müller

**Affiliations:** Department of Biology, École Normale Supérieure, PSL University, Paris, France; Institute of Computational Biology, Helmholtz Zentrum München, Munich, Germany; Department of Statistics, Ludwig-Maximilians-Universität München, Munich, Germany; Center for Computational Mathematics, Flatiron Institute, New York, USA

**Keywords:** irritable bowel syndrome, 16S rRNA amplicon data, meta-analysis, statistical analysis

## Abstract

**Background:** Irritable Bowel Syndrome (IBS) is a chronic functional bowel disorder causing abdominal discomfort, as well as transit deregulation with constipation and/or diarrhea. The pathophysiology of IBS is poorly understood and believed to be multifactorial. The role of gut microbiota in IBS has been investigated in several case-control studies, in particular via 16S rRNA amplicon sequencing surveys. These studies, however, have not yet led to a consistent picture of significant changes in gut microbial compositions across health and disease. One key bottleneck is the modest cohort sizes of most individual studies and a high diversity of experimental, bioinformatics, and statistical analysis approaches across studies.

**Results:** We address these shortcomings by presenting MetaIBS, an open-access data repository and associated meta-analysis workflow of thirteen 16S rRNA amplicon datasets comprising both fecal matter and sigmoid biopsy samples spanning ∽**2,500** IBS and healthy individuals. MetaIBS includes a tailored computational framework that (i) enables coherent *de novo* processing and taxonomic assignments of the raw 16S rRNA amplicon reads across experimental protocols and sequencing technologies, and (ii) statistical workflows for visualization and analysis at different taxonomic ranks and data granularity. Our statistical meta-analysis shows that popular high-level microbiome summary statistics, including Firmicutes/Bacteroidota ratios or diversity indices, are insufficient for reliable discrimination between IBS patients and healthy controls. Fine-grained multi-method differential abundance and classification analysis, however, can identify sets of differentially abundant taxa that replicate *across multiple datasets*, including *Coprococcus eutactus* and *Alistipes finegoldii*.

**Conclusions:** MetaIBS provides a curated and reproducible data and (meta-)analysis resource for amplicon-based IBS research at unprecedented scale. MetaIBS allows assessing the heterogeneity of IBS cohorts across multiple experimental protocols, sample types, and IBS phenotypes. Our framework will likely contribute to more coherent insights into the role of the microbiome in IBS and the discovery of reliable microbial IBS biomarkers for follow-up functional and translational studies.

## Background

Irritable Bowel Syndrome (IBS) is a chronic intestinal disorder affecting approximately 10-25% of the general population worldwide. It is characterized by (1) frequent abdominal pain, defined by one or more days per week for the previous three months; (2) an absence of alarming signs such as gastrointestinal bleeding, palpable abdominal mass or personal history that may suggest the presence of colorectal cancer (CRC) or inflammatory bowel disease (IBD); and (3) at least two out of these other symptoms: pain during defecation, change in stool frequency and/or change in stool morphology [1, 2, 3, 4, 5]. Its diagnosis relies on the Rome criteria (currently Rome IV), previously enumerated, defining it into four subtypes: constipation (IBS-C), diarrhea (IBS-D), mixed (IBS-M) or unclassified [1]. While IBS is a chronic syndrome impairing quality of life, it remains poorly understood with no curative treatment to date [1], thus requiring continued concerted scientific investigations into this peculiar idiopathic disease.

The roots of IBS pathophysiology are likely multifactorial, with reported alterations in gastrointestinal motility, visceral sensitivity, gut inflammation, and intestinal permeability [1, 4, 6]. All these elements can be modulated by the microbiome residing in the gut [6], and with increasing evidence of its role in other human diseases [7], IBS is strongly suspected to be related to some dysbiosis of the gut microbiota. Indeed, experimental systems with fecal transplantation of gut microbes from IBS patients to germ-free mice demonstrated alterations in the gut function, notably in the gastrointestinal transit, the permeability of the intestinal barrier, and the immune system homeostasis [8, 9]. In addition, acute gastroenteritis infections increase the risk of developing IBS by 4.2-fold in the year following the infection [10, 5], a phenomenon known as post-infectious IBS. Taken together, cumulative reports suggest a role of the gut microbiome in triggering and/or sustaining IBS symptoms. However, a defining gut microbial feature responsible for IBS has yet to be elucidated [1, 11, 12].

The gut microbiota is a complex ecosystem of various microorganisms, including yeast, protozoa, archaea, viruses, and predominantly bacteria [7]. Despite numerous studies comparing the gut microbiome composition in IBS patients and healthy controls, few of them reported consistent results [12, 11, 6]. Some recurrent evidence of dysbiosis include alterations in the Firmicutes to Bacteroidota (F/B) ratio [13, 14, 15, 16], and in the relative abundance of members of the Clostridiales order [14, 15, 2, 17, 18], and the *Ruminococcaceae* [14, 19, 17, 20, 21, 22, 23], *Streptococcaceae* [21, 15, 13, 24, 25], or *Enterobacteriaceae* [22, 23, 17, 26, 27] families. Although these microbial taxa are regularly highlighted in case-control studies, there is no consensus on whether they are increased or decreased in IBS [11, 12, 28].

These inconsistencies can be explained first and foremost by limited cohort sizes (many studies have 50 or fewer IBS patients), as well as the large heterogeneity in enrollment criteria, geographical localization, and thus, dietary habits. Moreover, differences in experimental protocols, amplified variable regions and sequencing technologies may have influenced the microbial species detected.

Finally, in the case of analysis of 16S rRNA sequencing data, studies have employed diverse bioinformatics pipelines (e.g., Qiime, Mothur, Usearch, Uparse) [29, 30, 31, 32], reference databases for the taxonomic alignment, such as, e.g., Silva, Greengenes, Ribosomal Database Project) [33, 34, 35, 36], and statistical analysis approaches, all of which may further contribute to inconsistencies. Therefore, comparisons between studies or interpretation of gut microbial alterations in IBS patients are limited.

To overcome the limitations of small cohort sizes and controversial findings due to varying protocols and bioinformatics pipelines, we introduce a standardized meta analysis of 16S rRNA sequencing data across thirteen IBS studies, called MetaIBS, enabling consistent comparison of gut microbiome compositions in IBS patients versus healthy controls. To this end, we inferred Amplicon Sequence Variants (ASVs) *de novo* from raw 16S rRNA read sequences using a standardized bioinformatic pipeline with a single reference database, thus enabling merging and cross-comparison of all datasets. All data files and associated fully documented and reproducible processing scripts are publicly available at https://github.com/bio-datascience/MetaIBS, thus providing a central resource for the scientific community to gain insights into the role of the gut microbiota in IBS pathophysiology. To illustrate the capabilities of this resource, we performed statistical analysis of a total of 2651 samples from 2356 individuals across ∼80,000 ASVs, using state-of-the-art log-ratio, α-diversity, dimensionality reduction, differential abundance, and regression analysis techniques.

We hypothesized that this approach would reduce inconsistencies across studies, and, ultimately, reveal insights on the role of the gut microbiome in IBS patients.

We found that high-level microbiome composition descriptors, such as F/B ratios or α-diversity, are inconsistent for IBS diagnosis. However, fine-grained multi-method differential abundance testing revealed 38 microbial genera that were differentially abundant in at least three datasets, most of which belonged to the *Ruminococcaceae* or *Lachnospiraceae* families (Clostridia class) and were decreased in IBS samples. MetaIBS also enabled cross-study identification of eight differentially abundant taxa on the strain level by merging two datasets that amplified the same variable region of the 16S rRNA gene and contained subsets of identical ASVs.

## Results

### Data collection and pre-processing

We collected datasets of 16S ribosomal RNA (16S rRNA) sequencing studies and analyzed the data with a standardized pipeline. We included published case-control studies that compared the microbiome composition of healthy and IBS individuals using 16S rRNA sequencing. Studies in children and interventional clinical studies were excluded. We identified 41 studies matching our inclusion criteria, as shown in Figure 1, thirteen of which either had their raw data publicly available or were kindly shared after inquiry by the authors [15]. Details on the included datasets are summarized in Table 1. We will refer to specific datasets by the name of the first author of the respective study (Table 1). Individuals were all diagnosed with IBS by a physician based on Rome III or Rome IV criteria. The Rome IV criteria have a stricter definition of abdominal pain frequency; however, 85% of patients with Rome III criteria also fulfill the Rome IV criteria [37]. From the American Gut Project (AGP) [38], we only included samples from individuals who reported being diagnosed by a medical professional (see Materials & Methods). Hence, we ensured that the IBS diagnosis criteria across datasets were comparable. In datasets with available covariates, we verified that age, BMI, and gender distributions were comparable between healthy and IBS individuals (Supplementary Table S1). The Lo Presti dataset [39] had significantly more women in the IBS group, and in the Zhu dataset [40], IBS individuals were significantly older than their healthy counterparts (Supplementary Table S1). In all other datasets with available covariates, cases and controls had comparable demographic characteristics.

**Figure 1.**
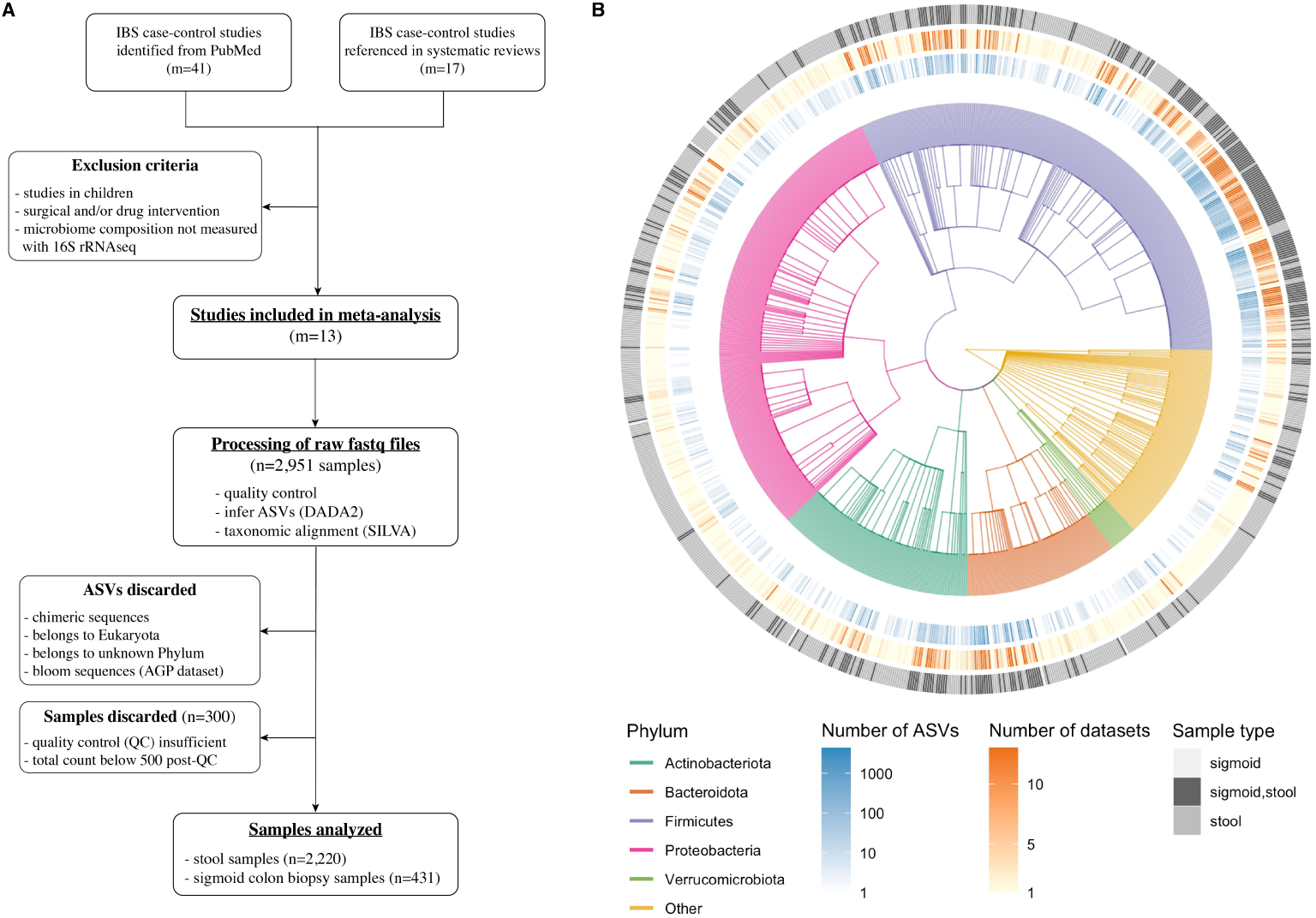
Overview of the meta-analysis. **(A)** Flowchart summarizing the identification of datasets that used 16S rRNA sequencing to compare microbiome composition in healthy versus IBS (case-control studies), then the re-processing of raw fastq files from the included datasets, followed by the final number of samples that passed the filtering steps. **(B)** Taxonomic tree representing the microbial genera detected in all samples. The outer circles represent the number of ASVs across all datasets that were found to belong to a specific genus (blue gradient); the number of datasets in which a specific genus was present (yellow gradient); and whether each genus was found in only in fecal samples, sigmoid colon biopsy samples, or both (grey scale).

**Table 1.** Characteristics of included datasets.

| Studies | Country | # samples analyzed (HC/IBS) | IBS diagnosis | DNA isolation protocol | Sequencing platform | Variable region (primers) | Reference |
| --- | --- | --- | --- | --- | --- | --- | --- |
| Labus et al., 2017 | USA | 23/29 | Rome III | PowerSoil Kit (MO BIO) | 454 pyrosequencing | V3-V5 (357F/926R) | [13] |
| Lo Presti et al., 2019 | Italy | 34/23 | Rome IV | DNA Stool Mini Kit (QIAGEN) | 454 pyrosequencing | V1-V3 (28F/519R) | [39] |
| Ringel-Kulka et al., 2015 | USA | 75 | Rome III | Mechanical disruption, phenol/chloroform extraction, alcohol precipitation | 454 pyrosequencing | V1-V2 (8F/357R) | [25] |
| AGP, 2021 | USA | 594/589 | N/A | PowerSoil Kit (QIAGEN) | Illumina MiSeq | V4 (515F/806R) | [38] |
| Liu et al., 2020 | China | 44/84 IBS-D | Rome IV | E.Z.N.A. Soil DNA Kit (Omega Bio-tek) | Illumina MiSeq | V3-V4 (338F/806R) | [41] |
| Pozuelo et al., 2015 | Spain | 88/185 | Rome III | Mechanical disruption with beads, alcohol precipitation | Illumina MiSeq | V4 (515F/806R) | [19] |
| Fukui et al., 2020 | Japan | 26/84 | Rome III | GENE PREP STAR PI-480 (Kurabo Industries) | Illumina MiSeq | V1-V2 (27F/338R) | [23] |
| Hugerth et al., 2020 | Sweden | 404/121 | Rome IV | ZR-96 Genomic DNA MagPrep (Zymo Research) | Illumina MiSeq | V3-V4 (341F/805R) | [28] |
| Mars et al., 2020 | USA | 24/45 | Rome III | PowerSoil Kit (QIAGEN) | Illumina MiSeq | V4 (515F/806R) | [24] |
| Zhu et al., 2019 | China | 14/15 | Rome III | PowerSoil Kit (MO BIO) | Illumina MiSeq | V4 (515F/806R) | [40] |
| Zhuang et al., 2018 | China | 10/20 IBS-D | Rome III | PowerFecal Kit (MO BIO) | Illumina MiSeq | V3-V4 (338F/806R) | [42] |
| Nagel et al., 2016 | Australia | 15/15 IBS-D | Rome III | DNA Stool Mini Kit (QIAGEN) | Ion Torrent | V4 (515F/806R) | [15] |
| Zeber-Lubecka et al., 2016 | Poland | 17/73 | Rome III | DNA Stool Mini Kit (QIAGEN) | Ion Torrent | Ion 16STM Metagenomics Kit (V2-4-8, V3-6, V7-9) | [14] |

MetaIBS comprises data from three sequencing technologies, including Illumina MiSeq [41, 19, 23, 28, 24, 40, 42, 38], 454 pyrosequencing [13, 39, 25], and Ion Torrent [15, 14] (see Table 1). For all samples, we inferred *de novo* amplicon sequence variants (ASVs) from raw FASTQ files and assigned taxonomy with the Silva reference database [33, 34]. More details on the standardized pre-processing pipeline and the number of samples that passed quality control can be found in Supplementary Tables S2 and S4, as well as the code repository at https://github.com/bio-datascience/MetaIBS.

In total, 79,943 ASVs were detected across 2,651 samples, with 2,220 stool samples and 431 biopsy samples obtained from the sigmoid colon mucosa (Fig. 1A). These ASVs belonged to 48 phyla and 973 known genera (see Fig. 1B), two thirds of which were detected in only one dataset. Genera were almost exclusively found either in stool samples only, or in both stool and sigmoid colon biopsy samples (Fig. 1B). Biopsy samples exhibited a higher proportion of Firmicutes (Supplementary Fig. S2B) compared to stool samples.

Overall, our standardized bioinformatics pipeline offers a coherent processing of raw 16S rRNA sequencing reads into ASV and taxonomic tables. This sets the stage for robust downstream statistical analyses to identify replicable microbial IBS biomarkers across datasets.

### Firmicutes to Bacteroidota ratios and α-diversity are not always altered in IBS

The literature on gut microbiome compositions in IBS gives inconclusive results regarding changes in high-level indicators such as α-diversity or changes in the ratios of the main bacterial phyla Firmicutes, Bacteroidota, or Proteobacteria [11]. We thus examined first whether standardized processing and analysis approaches allow to observe a consistent change in the commonly reported Firmicutes to Bacteroidota (F/B) ratio and/or α-diversity across the MetaIBS datasets.

To get a first overview of the data, we examined the relative abundances of the top five most abundant phyla across all samples (Supplementary Fig. S3). As expected, Firmicutes and Bacteroidota represented the majority of the gut microbial composition in both fecal and sigmoid colon biopsy samples, followed by Proteobacteria, Actinobacteriota, and Verrucomicrobiota (Supplementary Fig. S3A,B). Sigmoid colon biopsy samples exhibited an overall higher abundance of Firmicutes and lower abundance of Bacteroidota compared to fecal samples (Supplementary Fig. S3B,D). Despite healthy and IBS samples showing high variability in their phylum composition (Supplementary Fig. S3A), the average phylum composition in each disease group was stable across time points in both fecal or sigmoid colon biopsy samples, as seen in the Pozuelo and Mars datasets, respectively (Supplementary Fig. S3E). We did not observe any noticeable changes in the relative abundance of the main phyla between healthy and IBS samples (Supplementary Fig. S3D), which is partly explained by the high variability in observed phylum compositions (Supplementary Fig. S3A). The IBS- M subgroup did show higher abundance of Bacteroidota compared to healthy controls (Supplementary Fig. S3C). However, IBS subtypes were represented unequally across datasets, and the observed IBS-M microbiome compositions largely come from the Pozuelo and Zeber-Lubecka datasets and consist of fewer samples (n=67). Future datasets with more IBS-M samples are required to confirm the relevance of this observation.

To get a quantitative asssessment of potential alterations in the main phyla, we compared the F/B ratio between healthy and IBS individuals. In fecal samples, the F/B ratio was significantly increased in two datasets, while decreased in two other datasets (Fig. 2A). The majority of datasets had no significant change in the F/B ratio, both in fecal and sigmoid mucosa samples. In the Pozuelo dataset, we observed a significant decrease in F/B ratio in the IBS group compared to the healthy controls across both available time points (Fig. 2C). We next asked whether alterations in F/B ratio may be dependent on the IBS subtype (Fig. 2D). In fecal samples, we found a significant increase in F/B ratio in IBS-D patients from the Labus dataset, but observed the opposite for the Pozuelo data. Similarly, the Mars and Zeber-Lubecka datasets had a significant increase in F/B ratio in IBS-C patients, while the LoPresti dataset showed a significant decrease. However, since the number of samples in IBS subgroups is small (more than half of the subgroups have n < 15 samples), these observations may be skewed by outliers. In the AGP dataset, where IBS subtypes were presumed from available metadata on bowel movement quality and frequency, we observed no significant change in F/B ratio between healthy controls and IBS patients reporting diarrhea or constipation (Supplementary Fig. S5). Taken together, these observations show no consistent alteration in the F/B ratio across cohorts, despite using a standardized bioinformatic and statistical framework.

**Figure 2.**
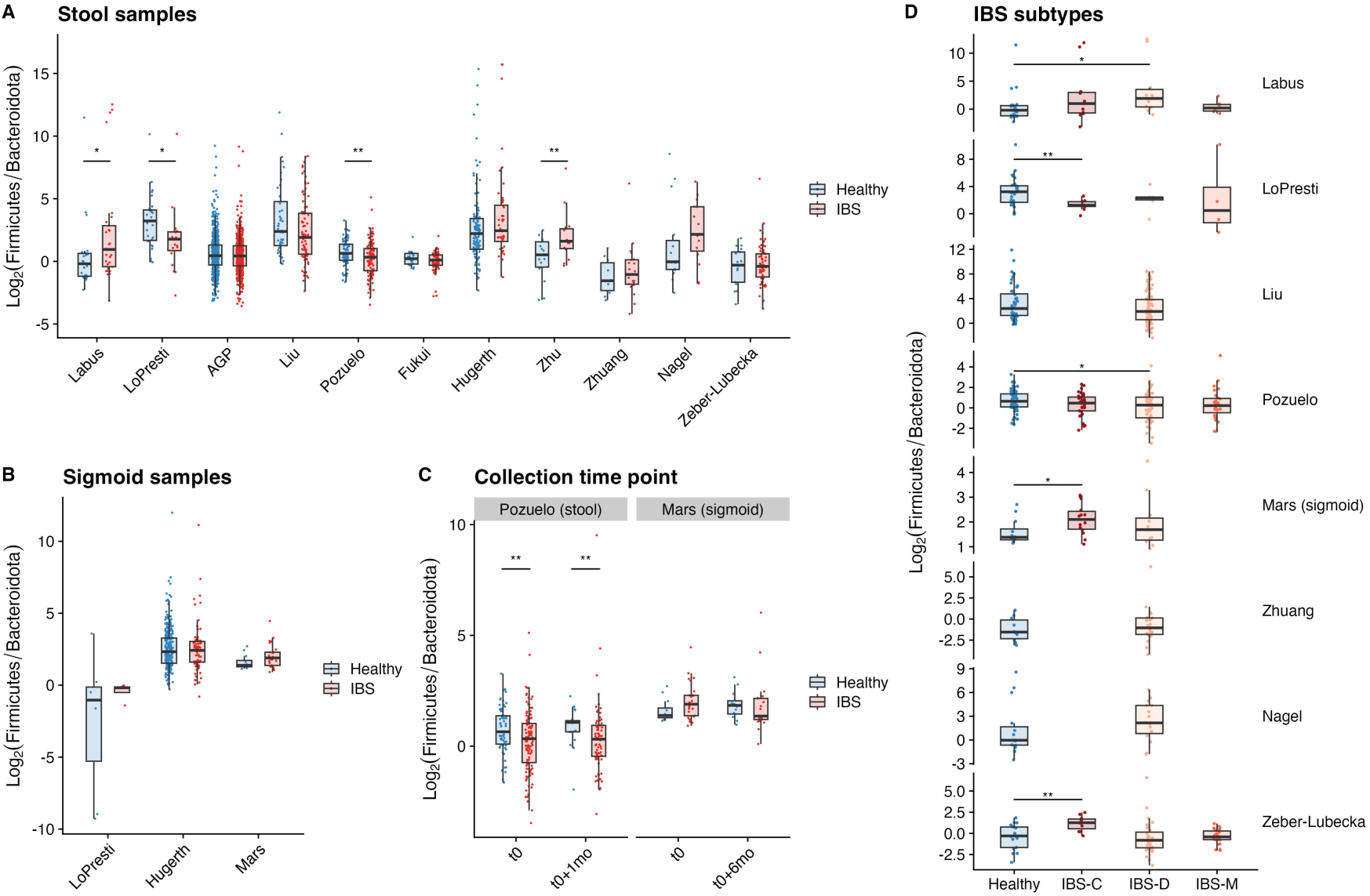
Log ratio of Firmicutes:Bacteroidota absolute count in **(A)** all fecal samples, as shown for each dataset separately (only 1st collection time point for Pozuelo); **(B)** all biopsies of sigmoid colon, as shown for each dataset separately (only 1st collection time point for Mars); **(C)** samples from a first or second collection time point in the Pozuelo and Mars datasets; **(D)** datasets with information on IBS subtype (constipation IBS-C, diarrhea IBS-D, mixed IBS-M). Statistical significance determined by a two-way wilcoxon test (* p*<*0.05; ** p*<*0.01; *** p*<*0.001).

Next, we asked whether the α-diversity was altered in IBS samples, a heavily debated topic in the literature [11, 12, 28]. In fecal samples, the Shannon index was significantly decreased in IBS patients in the Labus, Pozuelo, and Fukui datasets, but increased in the Zhu cohort (Supplementary Fig. S4A). This was also the case when measuring the Simpson index, with the exception of the Labus dataset, where we observed no significant change in α-diversity in IBS samples (Supplementary Fig. S4B). When looking more specifically at IBS subtypes, the Labus and Pozuelo datasets showed a significant decrease in α-diversity in IBS-C and in IBS-D patients, respectively, compared to healthy controls (Supplementary Fig. S4C). The majority of datasets showed no changes in the gut microbiome’s α-diversity between IBS patients and their healthy counterparts.

While previous works reported contradicting results on alterations of F/B ratio or α-diversity in IBS patients, our standardized pipeline still could not resolve these inconsistencies across cohorts. This suggests that these inconsistencies across studies are not due to the difference in processing and statistical methods, but may simply reflect that the F/B ratio and α- diversity are not defining features of IBS.

### Microbial composition is more affected by sequencing technology than IBS status

To assess whether there exist apparent global IBS “microbiome signatures” *across samples and datasets*, we next performed a host of unsupervised exploratory data analysis techniques including principle coordinate analysis on individual datasets, and clustered heatmap visualization and Uniform Manifold Approximation and Projection (UMAP) [43] on the entire MetaIBS corpus.

Since different variable regions of the 16S rRNA gene were amplified across studies, no *globally shared* sets of ASVs were available (see, however, Fig. 5A for subsets of shared ASVs). To enable cross-study analysis and prevent samples from clustering by dataset-specific ASVs, we first aggregated taxa to the family level. Figure 3A shows a heatmap of all microbial family abundances that were present in at least three MetaIBS datasets. We then computed pairwise Aitchison distances [44] (i.e., Euclidean distances between log-ratios of microbial family compositions) between all samples and constructed a low-dimensional embedding of the microbial compositions using UMAP [43] (Fig. 3B).

**Figure 3.**
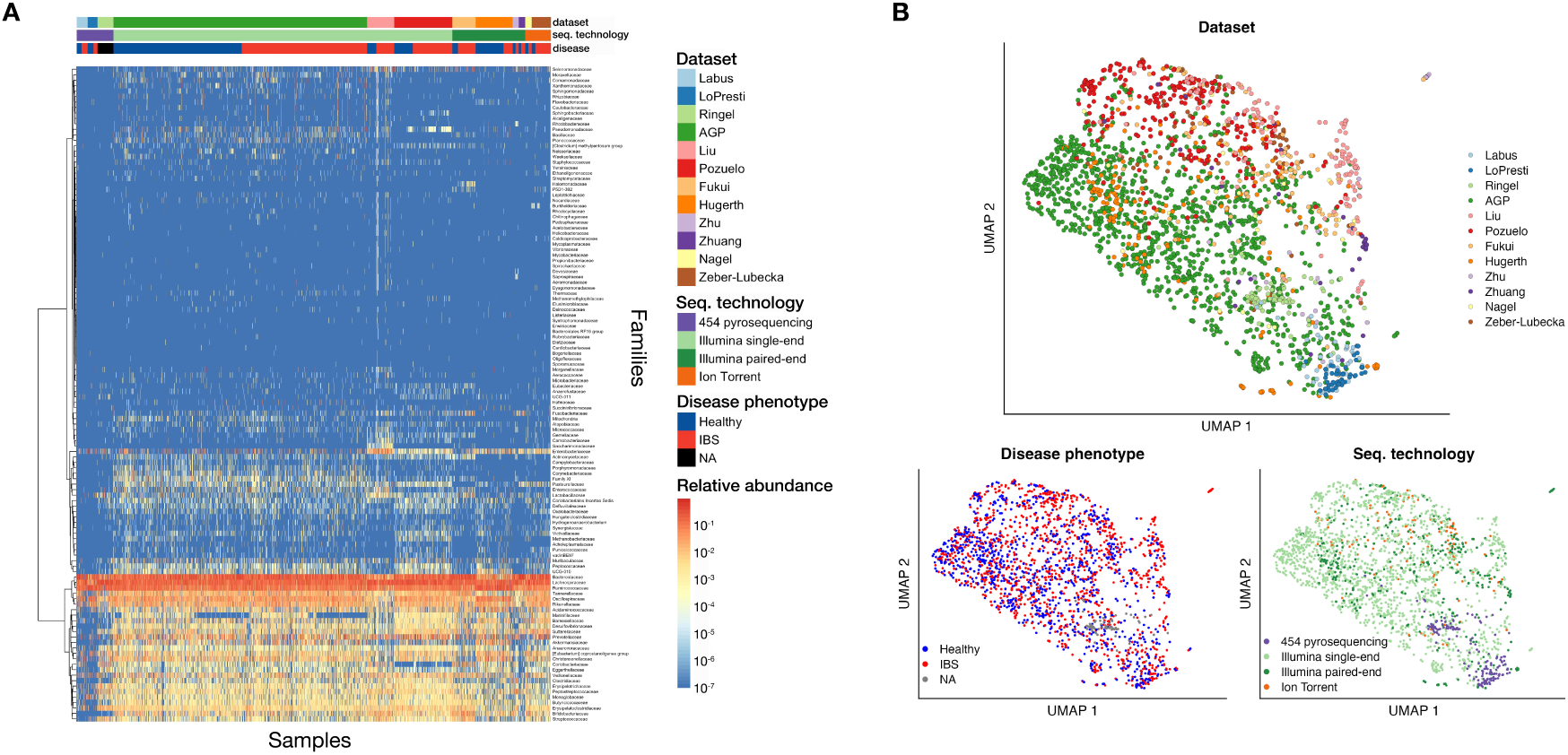
Sequencing technology biases microbial composition. **(A)** Heatmap of microbial families abundance in all fecal samples. Only families present in at least 3 datasets are shown. Families are in rows, clustered with the ward algorithm; samples are in columns, ordered by dataset and disease phenotype. Counts were normalized (sum per sample is 1). **(B)** UMAP was run on log-ratios between microbial families in all fecal samples. Samples are colored by dataset (top), disease phenotype (bottom left), or sequencing technology (bottom right). **(A-B)** All fecal samples were plotted (n=2,220), including both collection time points in the Pozuelo dataset.

The heatmap of family relative abundance patterns revealed a clear patterning by sequencing technology (in particular, 454 pyrosequencing vs others), and on a more fine-grained level, by dataset (e.g., Liu vs others) (Fig. 3A). Disease status, on the other hand, did not induce apparent large-scale abundance profile differences within individual studies. We confirmed this behavior by testing whether the compositional means of the abundant families were significantly different across datasets and across disease status, respectively, using pairwise compositional mean equivalence tests (see Methods)[45]. We confirmed that mean compositions across all pairs of datasets were significantly different whereas, with respect to IBS status, only the Zhu, the Pozuelo, and the Fukui datasets showed significant changes in compositional means.

Similarly, sigmoid colon biopsy samples showed no clear distinction in microbial families between healthy and IBS individuals, and samples clustered partially by dataset (Supplementary Fig. S6A,B). This is also confirmed in the UMAP visualization. Whereas samples from the American Gut Project (AGP) cover a large extent of the UMAP, all other datasets span distinct regions in the map (Fig. 3B, top panel). We observed no clear separation of IBS samples from the healthy ones in the low-dimensional projections (Fig. 3B, lower left panel), even when looking more specifically at IBS subtypes (Supplementary Fig. S6C). Rather, samples obtained by 454 pyrosequencing clustered separately from other sequencing technologies (Fig. 3B, lower right panel). This suggests that, despite using the same denoising scheme and reference database for taxonomic alignment of ASVs in all datasets, observed microbial compositions are still highly dependent on the experimental protocol and dataset, and that this effect is greater than the healthy/IBS status of the host.

In order to eliminate experimental biases, we repeated our analysis in individual datasets with large cohorts, notably AGP, Hugerth, and Pozuelo. Since we conducted the analysis separately in each dataset, we did not need to aggregate taxa. We computed the Bray-Curtis dissimilarity and visualized samples by Principle Coordinate Analysis (PCoA) [46]. Once again, healthy samples overlapped with IBS samples (Supplementary Fig. S7). Interestingly, in the Pozuelo dataset, fecal samples from the same individual taken within a month’s interval showed large dissimilarities, suggesting higher variability at lower taxonomic levels compared to the phylum composition (Supplementary Fig. S3E). As expected, the first axis separates fecal and sigmoid colon biopsy samples in the Hugerth dataset (Supplementary Fig. S7). Overall, this suggests that IBS patients do not exhibit major alterations in their microbial communities, and that there exist no broad microbial signatures associated with IBS.

Taken together, the present exploratory data analysis revealed that (i) experimental protocols and sequencing technology have a larger effect on the observed microbial compositions than a host’s pathophysiology and that (ii) exploratory data analysis of individual datasets with larger cohort sizes did not reveal apparent shifts in community composition of IBS patients.

### Differential abundance testing reveals taxa associated with IBS across datasets

We next investigated whether we could consistently detect compositional shifts in subsets of microbial taxa in IBS patients across datasets. Specifically, we used three distinct state-of-the-art statistical methods, ANCOM-BC [47], LinDA [48], and scCODA [49] to find differentially abundant (DA) taxa across the eleven datasets containing stool samples (Fig. 4A/B, Supplementary Fig. S8).

**Figure 4.**
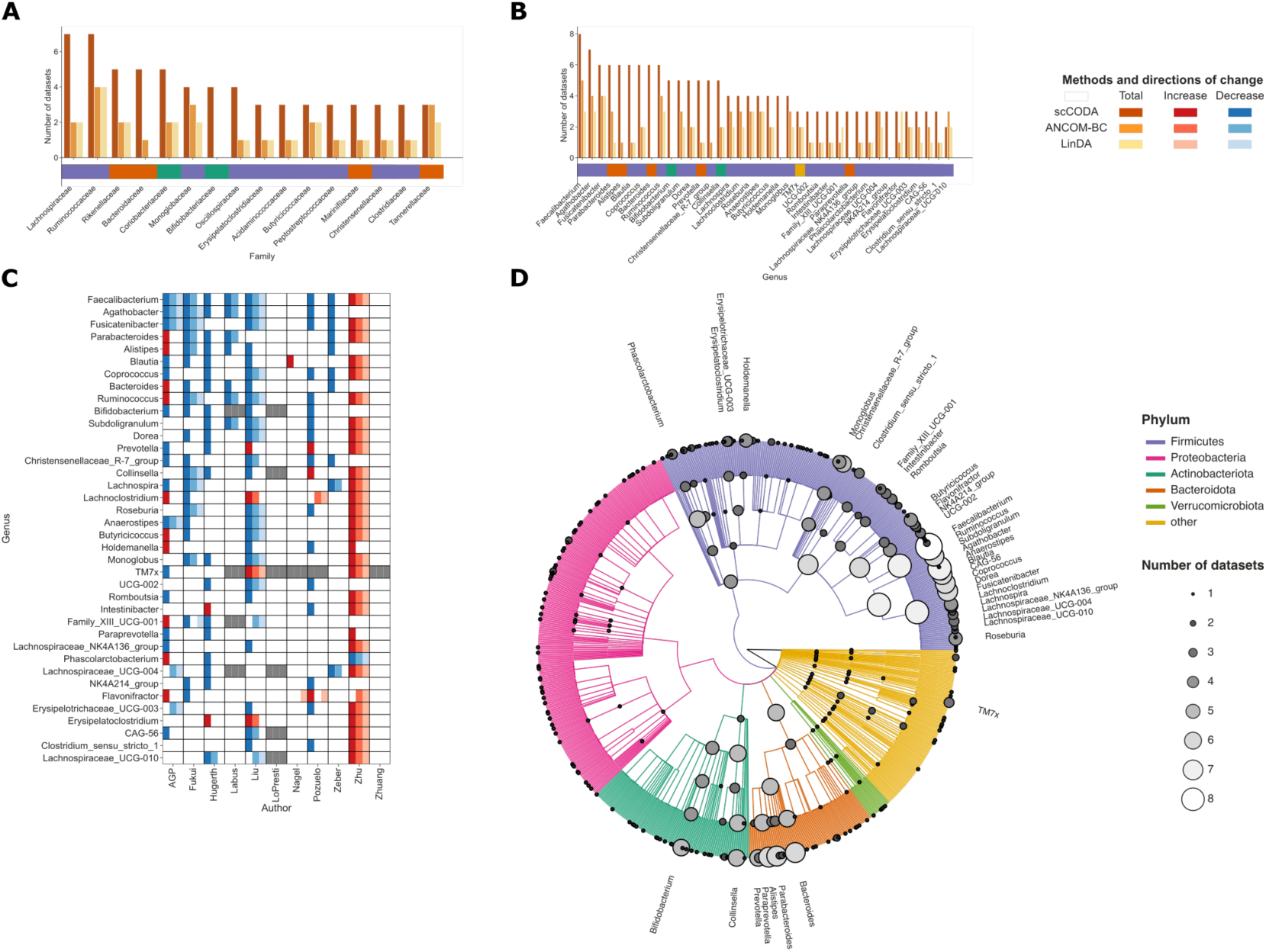
Differential abundance testing with three different methods reveals common trends in stool samples across 11 different datasets. **(A-B)** Aggregation to family and genus levels shows consistency of differential abundance in subpopulations of the gut microbiome between healthy individuals and IBS patients. Only families/genera that were determined as differentially abundant by at least one method in at least three datasets are shown. Colored boxes below the x-axis show the corresponding phylum for each family. **(C)** Heatmap of differential abundances on the genus level. Red boxes indicate a significant or credible increase for all three methods (scCODA, ANCOM-BC, LinDA), blue boxes a decrease. Grey fields show that the genus was not present in any sample of the respective dataset. The selection of genera is equal to the ones shown in (B). **(D)** Taxonomic tree including all genera that are present in at least one dataset (stool samples only). Grey circles on the nodes and leaves indicate the number of datasets that a taxon was differentially abundant in, as determined by scCODA. The branch colors indicate the phylum, tip labels denote the 38 genera from (B) that were found to be differentially abundant in at least three datasets.

We analyzed taxa on phylum, class, order, family, and genus rank, respectively, and recorded both how often the different methods agreed on DA taxa and how often DA taxa were identified across datasets. Overall, scCODA identified the largest number of DA taxa irrespective of taxonomic ranks. In general, all methods identified more DA taxa on lower taxonomic ranks. Figure 4D visualizes all DA taxa found by scCODA on the taxonomic tree and reports how often a particular DA taxon was identified across the eleven datasets. On the family level (Fig. 4A), *Ruminococcaceae* and *Lachnospiraceae* were most often found to change in relative abundance, with scCODA finding a credible change in seven datasets, while ANCOM-BC and LinDA determined a significant shift four and two times, respectively. Furthermore, ten out of the 16 families that scCODA determined to be differentially abundant in more than two datasets belonged to the Firmicutes phylum.

On the genus level, Firmicutes made up an even larger share of DA taxa found in at least three datasets (30 out of 38, Fig. 4B). We considered these taxa as “genera of interest”, as changes in their abundance could possibly be associated with IBS. *Faecalibacterium* was the most consistently detected DA genus where a change in relative abundance was detected in eight datasets by scCODA, in five datasets by ANCOM-BC, and in three datasets by LinDA, respectively. This is consistent with multiple other studies [50, 17, 23] that also observed changes in *Faecalibacterium* abundance. DA genera that were detected in more than half of the datasets (by any of the methods scCODA/ANCOM-BC/LinDA) include *Agathobacter* (7/4/3 DA datasets), *Fusicatenibacter* (6/4/4 DA datasets), *Ruminococcus* (6/4/3 DA datasets), *Parabacteroides* (6/3/1 DA datasets), *Blautia* (6/1/1 DA datasets), *Coprococcus* (6/2/2 DA datasets), *Alistipes* (6/1/0 DA datasets), and *Bacteroides* (6/0/0 DA datasets), respectively (see also Fig. 4B). It is noteworthy that many of the identified DA genera were also found in the original analyses of the respective datasets as well as other IBS studies [11, 23, 51].

Next, we analyzed the direction of change (relative increase or decrease) in each dataset for the 38 genera of interest from Figure 4B. Figure 4C summarizes the direction of change across all methods and datasets. We observed that the majority of genera were found to decrease in relative abundance in IBS samples (indicated in blue in Fig. 4C). In particular, the genera *Faecalibacterium*, *Agathobacter*, and *Fusicatenibacter* showed remarkable consistency in direction of change across methods and datasets.

The genera *Flavonifractor*, *Lachnoclostridium*, *Intestinibacter*, *Prevotella* and *TM7x* showed increased relative abundance in IBS for a majority of datasets (indicated in red, in figure 4C). The dataset from Zhu et al. [40] appears as an outlier, since all three methods determined many genera to increase in relative abundance in IBS patients compared to healthy controls, contrary to a significant decrease in other datasets. This observation likely stems from a sequencing bias, discovered in the pre-processing of the raw 16S rRNA sequencing data where the forward primer (F515) was present at the end of reverse reads in IBS samples, but not in healthy samples. This led to *disjoint* sets of ASVs in healthy and IBS samples, and this bias persisted in higher taxonomic ranks.

In general, the number of DA genera per dataset varied considerably, as summarized in Table 2. Two datasets (Lo Presti et al. [39], Zhuang et al. [42]) contained no DA taxa regardless of the statistical method, while almost half of all genera were differentially abundant when analyzing the data from Zhu et al. using LinDA.

**Table 2.** Total number of genera, as well as number and share of differentially abundant genera, as determined by three different DA testing methods on each dataset (only stool samples).

| First author | Total genera | DA genera (fraction) |  |  |
| --- | --- | --- | --- | --- |
|  |  | ANCOM-BC | LinDA | scCODA |
| Labus | 91 | 5 (0.055) | 0 (0.000) | 7 (0.077) |
| LoPresti | 90 | 0 (0.000) | 0 (0.000) | 0 (0.000) |
| AGP | 620 | 16 (0.026) | 20 (0.032) | 197 (0.318) |
| Liu | 698 | 105 (0.150) | 42 (0.060) | 54 (0.077) |
| Pozuelo | 309 | 4 (0.013) | 6 (0.019) | 34 (0.110) |
| Fukui | 210 | 14 (0.067) | 10 (0.048) | 31 (0.148) |
| Hugerth | 263 | 2 (0.008) | 0 (0.000) | 31 (0.118) |
| Zhu | 143 | 59 (0.413) | 71 (0.497) | 41 (0.287) |
| Zhuang | 116 | 0 (0.000) | 0 (0.000) | 0 (0.000) |
| Nagel | 163 | 3 (0.018) | 7 (0.043) | 1 (0.006) |
| Zeber-Lubecka | 236 | 4 (0.017) | 1 (0.004) | 13 (0.055) |

Finally, we compared how the differentially abundant taxa (as determined by scCODA) are related in terms of taxonomy (Fig. 4D). Notably, most taxa that were differentially abundant in six or more datasets are genera that are parts of the *Firmicutes Clostridia Lachnospirales Lachnospiraceae* and *Firmicutes Clostridia Oscillospirales Ruminococcaceae* families, or higher order taxonomic groups that contain these. We noticed the same trend, although less clear due to the number of differentially abundant taxa being generally lower, also for ANCOM-BC and LinDA (Fig. S9).

In summary, while previous works often reported the microbial signature of IBS to be unclear and dependent on the cohort [11], our meta-analysis approach with standardized data processing and differential abundance analysis methods was able to determine 38 genera whose change of abundance was associated with IBS in multiple datasets with quantifiable degree of consistency.

### Analysis of shared ASVs between datasets can potentially reduce study-specific noise

The standardized preprocessing steps of the 16S rRNA sequencing data enabled combining observations from different studies. Focusing on the subset of shared ASVs between various studies could lead to more robust biomarker identification by reducing possible study-specific noise. Eight ASVs were pinpointed by all methods using classification analysis and differential abundance testing of shared ASVs between the Nagel [15] and Pozuelo [19] datasets. The methods were sparse log-contrast modeling for classification, LinDA, scCODA, and ANCOM-BC. The taxonomic assignment of the selected ASVs can be found in Figure 5D. Seven of these detected ASVs belong to the phylum Firmicutes, while one belongs to Bacteroidota. It is also noticeable that six of these ASVs belong to the class Clostridia. The taxonomic tree of the shared ASVs shows that Firmicutes and Clostridia build a large part of the shared ASVs (Fig. 5C). For three ASVs, we have a species-level annotation, namely *eutactus*, *AC2044*, and *finegoldii*.

**Figure 5.**
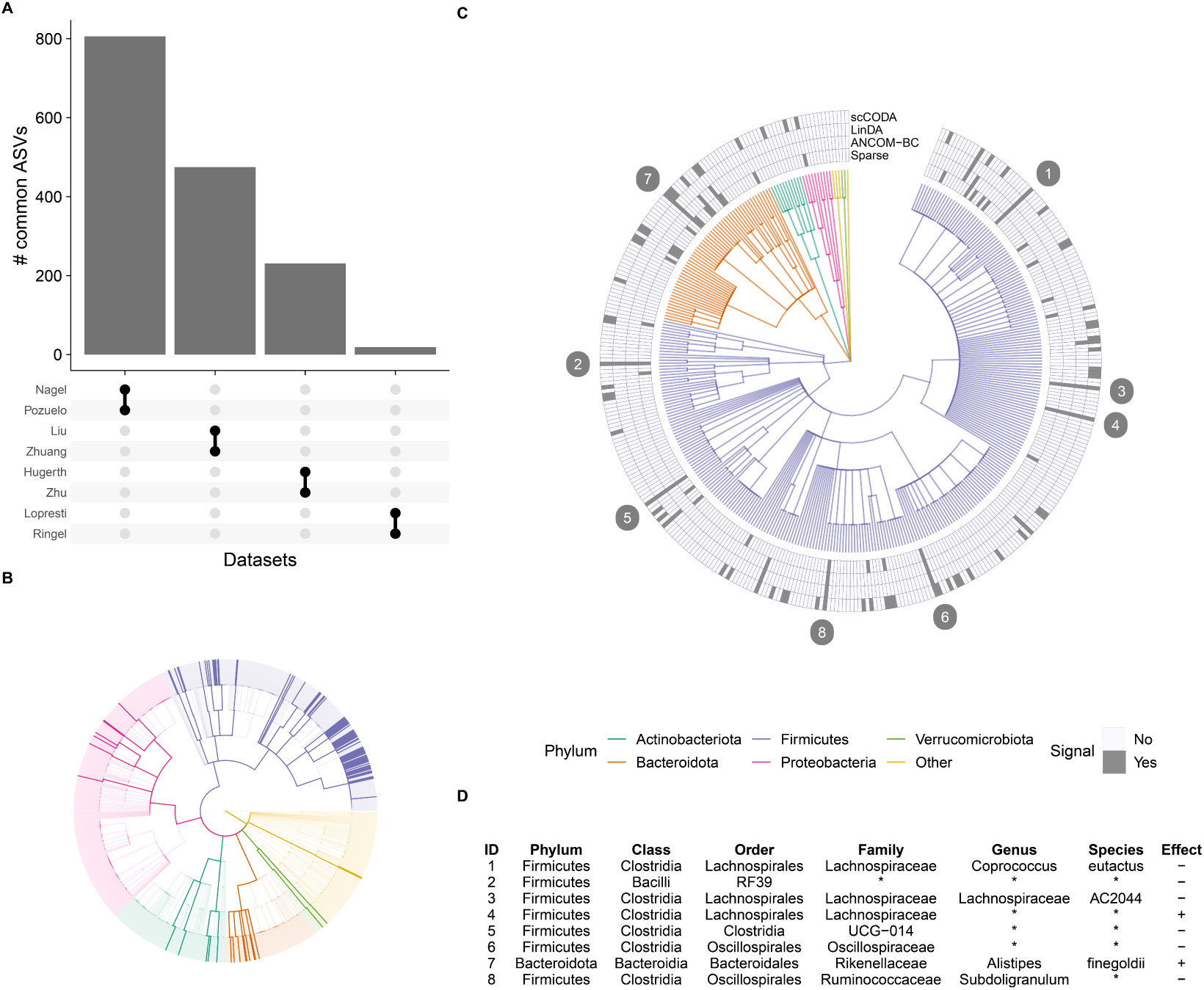
Analysis of shared ASVs. **(A)** Bar chart showing how many ASVs are shared by the different datasets. **(B)** Taxonomic tree displaying which genera are shared between the Nagel and Pozuelo datasets. The highlighted branches are present in the shared ASV subset of Nagel and Pozuelo. **(C)** Taxonomic tree of ASVs present in the Nagel and Pozuelo subsets. The outer ring indicates whether the method detects the ASV as differentially abundant or whether it was selected for classification by the sparse log contrast model. **(D)** The table reveals the taxonomic assignment for ASVs with signals detected by all four approaches. The ID is illustrated by the number in the circle next to the ASV of subplot (C). The three differential abundance and the classification methods share the same effect direction shown in the column effect. For the differential abundance methods, the plus indicates an increase, and the minus means a decrease in abundance. The sign suggests a positive or negative coefficient for the classification method.

Next to the 38 genera identified by differential abundance testing on the genus level, the eight identified ASVs shared between the Nagel and Pozuelo datasets can serve as a good starting point for further research.

## Discussion

Here we developed a framework to combine 16S rRNA sequencing data of IBS and healthy gut microbiota, with the purpose of standardizing bioinformatic preprocessing and statistical analyses for more robust and interpretable results on microbial alterations potentially occurring in IBS. By combining 13 datasets, we found that high-level analyses such as F/B ratio, α-diversity, or dimensionality reduction do not show significant *and* consistent changes across datasets. However, more fine-grained analysis looking at differentially abundant taxa at lower taxonomic levels showed that 38 genera were DA in at least three datasets, most of which belonged to the Firmicutes phylum and were significantly decreased in IBS patients. This observation was consistent using three well recognized statistical methods (ANCOM-BC, LinDA and scCODA) that take into account the compositional nature of microbiome data, which is paramount for correct statistical analysis [52]. Furthermore, our standardized meta-analysis allowed to identify ASVs shared across datasets that had amplified the same variable region of the 16S rRNA gene. Restricting our DA testing on these shared ASVs between the Nagel and Pozuelo datasets, we found eight ASVs that were consistently recognized as DA by the different statistical methods used, six of which belonged to the Clostridia class from the Firmicutes phylum, and most of which were significantly decreased in IBS.

Literature reports conflicting results on alterations in microbiome composition in IBS, every or most studies suggesting to have found a ”microbiome signature” in IBS [11, 12], either through an alteration of the F/B ratio [13, 53, 2, 15, 19, 42], α-diversity [13, 19, 40], and/or a set of specific microbes [40, 13, 2, 24, 41, 23, 19, 39]. However, more recent studies with bigger cohort sizes seem to suggest otherwise, with little to no microbiome signature between IBS and healthy controls, due to the high heterogeneity in microbiome composition in IBS patients [28, 54]. Our results confirm these findings, as we found no consistent alterations in F/B ratio or in α-diversity, despite using the same bioinformatic pipeline to preprocess the data and the same reference database for taxonomic alignment. These inconsistencies can be due to a variety of factors, including the small cohort sizes of most studies; the differences in experimental protocols (sample handling, DNA extraction, variable region amplified, sequencing technology); study localization and thus dietary habits of the participants; and last but not least, the inclusion criteria and demographics of the participants (BMI, age, sex, comorbidities, recent treatments, etc.). To note, the Pozuelo dataset is one of the biggest cohort sizes (113 IBS patients and 66 healthy controls) in our meta-analysis and in the literature [19], and we did see a significant reduction in the F/B ratio in that dataset, as was reported by the authors [19]. This alteration was stable with time, as one month later there was still a decrease in the F/B ratio in the same participants (Fig.2C), suggesting that this was not a one-time observation but potentially a long-lasting change in the microbiome of these patients. However, we did not observe any changes in F/B ratio in the Hugerth or AGP data, two other datasets with higher cohort sizes [28, 38]. Age and BMI are known to be factors influencing overall microbiome composition [55, 28], yet participants in the Pozuelo, Hugerth and AGP datasets have comparable BMI and age distribution (Supplementary Table S1, [19]). This discrepancy in the F/B ratio is thus probably explained by differences in experimental protocols, study localization, and other patient characteristics.

Similarly, we observed no consistent changes in α-diversity across datasets. Previous reports suggest that IBS patients have either a decreased or unaltered α-diversity compared to healthy controls [12], which overall aligns with our observations (Supplementary Fig. S4). Only IBS patients from the Zhu dataset had an increased α-diversity compared to their healthy counterparts, as reported by the authors [40], however in this dataset we observed that ASVs inferred from healthy and IBS samples were entirely distinct, which probably biased the microbial taxa detected and thus the α-diversity measurement. Surprisingly, we found that the Shannon index was decreased in IBS patients from the Labus dataset compared to healthy controls (Supplementary Fig. S4), whereas the original study reported the opposite finding [13]. To note, in that paper the authors split IBS samples into ”IBS” and ”healthy-like IBS” categories based on β-diversity analyses, and they showed that ”IBS” samples had higher α-diversity compared to ”healthy-like IBS” samples or healthy samples. This differs from our method, where we compared *all* IBS samples to healthy ones, and we observed only a small significant decrease in the Shannon index, or no changes in the Simpson index (Supplementary Fig. S4). We thus think that the contradiction in our observations from the same data may stem from the difference in the comparison groups, but also probably from the different preprocessing pipelines used (DADA2 vs QIIME, Silva vs Greengenes). Overall, we found that changes in α-diversity are not a defining feature of IBS, which is consistent with reports from recent studies with bigger cohort sizes [56, 54, 28], and also with the fact that it is not a reliable indicator of dysbiosis in other gut-related diseases [57]. This implies that the total number and the evenness of microbial species remains unchanged in IBS, however it does not provide any information on potential changes in the actual composition of the microbiome.

To that end, we compared the microbiome composition across samples by looking at the relative abundances of microbial families, and observed that samples were discriminated more by a study effect than by disease phenotype (Fig. 3, Supplementary Fig. S6). We expected that by aggregating data at the genus or family level, we would lose fine-grained analyses on microbial species while limiting batch effects, as has been done in a previous meta-analysis [57]. However, we still observed significant study-specific variations, even among microbial families. Despite including 13 datasets in this meta-analysis, there are still three features of experimental variability distinguishing studies: the DNA extraction protocol, PCR primers and sequencing technology, all of which have been shown to influence greatly the microbial taxa detected [55, 58]. Only the Zhu and AGP datasets have these three characteristics almost identical (Table 1), except that (1) samples were shipped by participants to the study center in the AGP dataset; and (2) single-end vs paired-end sequencing was performed in the AGP vs Zhu datasets respectively [38, 40]. Unfortunately, there are currently few options to correct for such batch effects in 16S rRNA sequencing [59]. In the future, extending this meta-analysis by adding more datasets, and also having new studies with standardized protocols and sharing their raw data, will help overcome this limitation. In the meantime, we observed that the microbial families present in fecal or sigmoid biopsy samples did not allow to discriminate healthy from IBS individuals, even by IBS subtype (Supplementary Fig. S6), or even by focusing on single datasets with the biggest cohort sizes (Supplementary Fig. S7). This result is coherent with β-diversity analyses in recent studies that have enrolled more subjects than average [28, 56, 54], where they also reported both in fecal and colonic biopsy samples that IBS patients did not exhibit a distinctive variation in their microbial communities.

Nevertheless, investigating more finely differential abundant taxa between cases and controls, we found 38 genera were DA in at least three datasets, including nine that were DA in six datasets or more (Fig. 4). The vast majority of these genera of interest belonged either to the *Ruminococcaceae* or *Lachnospiraceae* families, both of which are butyrate-producing Clostridia. Butyrate and other short-chain fatty acids (SCFAs) produced by these microbial families are not only essential nutrients for colonic epithelial cells [60, 61], but they also promote the accumulation and differentiation of regulatory T cells in the gut epithelium [60, 62], which are important to restrain inflammatory responses. The loss of such beneficial microbes may thus prevent proper function of the gut epithelial barrier. The Pozuelo study did report a decrease in *Ruminococcaceae* particularly in IBS-D patients, and hypothesized that this reduction could alter the gut epithelium permeability and contribute to the diarrhea [19]. This hypothesis goes along with the observation that higher relative abundance of *Ruminococcaceae* in fecal samples correlates with harder stools on the Bristol Stool Scale [63]. Considering that there is controversy on whether *Ruminococcaceae* is increased, decreased, or unaltered in IBS [11, 12], it is thus new that we find here multiple genera that are consistently less abundant in IBS samples compared to healthy controls in several datasets. To note, in our meta-analysis we may have an over-representation of IBS-D samples (n=277) compared to other subtypes (n=195), although there is a substantial number of samples of unknown IBS subtype (n=811), so the significant decrease in *Ruminococcaceae* that we observed may be biased towards IBS-D samples. Extending this meta-analysis with more datasets, especially with covariates on stool consistency and IBS subtype, will be paramount to confirm the association of *Ruminococcaceae* and *Lachnospiraceae* with IBS or with more specific IBS subtype(s).

In this meta-analysis, we were unfortunately unable to include many datasets derived from gut mucosal biopsy samples and thus conduct detailed statistical analyses on these samples despite their relevance. The microbial composition in fecal and sigmoid biopsy samples was drastically different, as expected [28, 24, 39], hence samples from the gut mucosa will better reflect the microbial ecosystem interacting with the gut epithelium. To overcome this limitation, we are making this meta-analysis publicly available to the scientific community, so that anyone can add their own dataset(s) and further investigate microbial alterations occurring in IBS. We also acknowledge that we report here only associations between IBS and microbiome composition, which is not proof of causation and needs to be confirmed experimentally.

In summary, our meta-analysis confirms that Firmicutes to Bacteroidota ratio and α-diversity changes are not defining features of IBS. Furthermore, strong study-specific effects urges the need to have established standardized protocols in 16S rRNA sequencing for comparable and reproducible research in the future. Finally, we identified multiple genera from the *Ruminococcaceae* and *Lachnospiraceae* bacterial families as being downregulated in fecal samples from IBS patients across several studies, which is the first time that such an alteration associated with IBS is consistently observed in multiple studies.

## Materials and methods

### Search strategy and data collection

Case-control studies comparing gut microbiome composition between IBS patients and healthy controls using 16S rRNA sequencing were first identified from references listed in systematic reviews [12, 11]. We extended our search further to PubMed using keywords such as ”irritable bowel syndrome” and ”gut microbiota”. We included in our meta-analysis any study published until April 2021 that had (1) enrolled adult IBS patients and healthy controls; (2) performed no surgical or drug intervention on patients in an effort to treat IBS; (3) measured gut microbiome composition with 16S rRNA sequencing. We obtained raw fastq files and sample covariates of the included studies either from online repositories (Sequence Read Archive, European Nucleotide Archive), or from personal communication with the authors [15]. Data from the American Gut Project (AGP) was also included in our meta-analysis [38]. As the AGP is an open platform with thousands of microbiome samples, we first downloaded the covariates table from the sequence read archive (SRA) on June 6th 2021, in order to identify samples of interest. The questionnaire filled in by the participants allowed us to discard any sample with (1) a reported age outside 18 - 70 years old; (2) a BMI outside 16 - 35; (3) any antibiotics taken in the past 6 months; (4) other reported comorbidities such as inflammatory bowel disease, *C.difficile* infection, fungal overgrowth in their gut, gluten intolerance or diabetes. This population consisted of 4,722 healthy samples and 645 samples diagnosed with IBS by a medical professional. We randomly chose 645 healthy samples to have an even number of cases and controls, verifying by a Student’s t-test that healthy and IBS samples had comparable age and BMI distribution (Supplementary Table S1). We thus downloaded a total of 1,290 fastq files from the AGP (Supplementary Table S4). The code for this sample selection strategy is available on our github repository (https://github.com/bio-datascience/MetaIBS).

### Processing of 16S rRNA sequencing data

For datasets that required demultiplexing (Supplementary Table S2), reads were assigned to each sample based on their barcode, and individual fastq files for each sample were generated using sabre (https://github.com/najoshi/sabre). We re-processed raw fastq files of 16S rRNA sequence data in R (v. 4.0.4) [64] using the DADA2 package version 1.21.0 [65], in order to perform quality filtering and obtain the count and taxonomic dataframes. These steps are further described hereafter. First, we discarded reads not containing primers, and trimmed off the primers and any preceding sequence.

To note, no primers were found in the AGP dataset, and in the Zeber-Lubecka dataset an unknown mix of primers was used, so this step was skipped (Supplementary Table S2). Then, we filtered reads based on their quality scores with the *filterAndTrim* function from DADA2 [65], using default parameters except that reads were (1) truncated at the first base where a quality score of 10 or below was observed; (2) discarded if shorter than 150bp; (3) discarded if they contained more than 3 expected errors. Next, error rates were estimated from randomly chosen samples with the *learnError* function [65]. We checked that the estimation from the parametric error model matched the observed error rates. For samples obtained from Ion Torrent sequencing technology, manual model adjustment was necessary at high quality scores to match the observed error rates (Supplementary Table S2). Subsequently, we identified *de novo* amplicon sequence variants (ASVs) with the *dada* function, which leverages the error model to assess whether sequence variants are more likely to be ASVs or amplicon errors [65]. For paired-end sequencing data, we merged paired reads after this step. After inferring ASVs, we looked at their length distribution and compared it to the supposed length of the amplified 16S variable region. This allowed us to discard abnormally short or long ASVs (50bp above or below the expected length), which are likely from non-specific priming. Chimeric sequences were also identified and discarded. Finally, ASVs were aligned to the Silva reference database (release 138) from the Kingdom to the Species level [33, 34]. After taxonomic alignment, we removed ASVs belonging to Eukaryota or to an unknown phylum. In addition, we discarded samples with a total count lower than 500. In the AGP dataset, as study participants ship their fecal samples to the nearest study center, certain bacterial taxa grow at room temperature and bias the observed microbiome composition [66]. Amir et al. [66] shared a list of operational taxonomic units known to bloom at room temperature (https://github.com/knightlab-analyses/bloom-analyses), which are advised to be removed when analyzing the AGP data. We thus discarded any ASVs identified as bloom sequences in the AGP dataset. Finally, we combined the ASV, taxonomic, and metadata tables into a phyloseq object using the Phyloseq package (v.1.34.0) for further analysis [67].

### Taxonomic tree

We combined phyloseq objects from all datasets with the *merge phyloseq* function from the Phyloseq package. To plot a taxonomic tree representing all detected genera across datasets, we first aggregated taxa to the genus level, before transforming the taxonomic table into a treedata object with the treeio pacakge (v.1.18.1) [68]. We then plotted the taxonomic tree with the ggtree package (v.3.2.1) [69].

### Heatmap

To plot a heatmap of the main microbial families represented among fecal samples, we aggregated taxa to the family level and divided counts by the total count per sample in order to obtain relative abundances. We then kept only families present in at least three datasets (p=116 families). We used the pheatmap package (v.1.0.12.; https://CRAN.R-project.org/package=pheatmap) to plot microbial families as rows, clustered using the Ward error sum of squares hierarchical clustering method with the method = ”ward.D2” option [70, 71]; and samples as columns, ordered by disease phenotype and dataset. For visualization purposes, we plotted the relative abundances on the heatmap on a log10 color scale.

### Uniform Manifold Approximation and Projection

To perform a Uniform Manifold Approximation and Projection (UMAP) [43], we used pairwise logratios between microbial families as input data. To obtain these, we aggregated taxa to the family level, then added pseudocounts of 0.5 before computing pairwise log-ratios between all microbial families across datasets. We computed the UMAP on all fecal samples (n=2,220) with the *umap* function from the uwot package (v.0.1.11.; https://github.com/jlmelville/uwot), using 20 neighbors and 3 components.

### Firmicutes to Bacteroidota ratio

In order to compute the Firmicutes to Bacteroidota (F/B) ratio, we first aggregated taxa to the phylum level in order to obtain the total count of each phylum per sample. We imputed a 0.5 pseudocount in samples containing no Firmicutes and/or Bacteroidota. Then, we calculated the log2 ratio of absolute Firmicutes over Bacteroidota counts in each sample. For datasets with two time points we only included samples from the first time point, while for datasets with both fecal and sigmoid biopsy samples we plotted the F/B ratio separately for each sample type.

### Sequencing depth

To visualize sequencing depth differences between datasets and the number of reads discarded through our standardized preprocessing pipeline, we plotted the number of reads before and after quality control of the data (Supplementary Fig. S1). We called the number of reads in the raw fastq files ”before” quality control, and the number of counts per sample in the ASV table right after removing chimeric sequences (see *Processing of 16S rRNA sequencing data*) ”after” quality control. Thus, this doesn’t take into account that we removed samples with less than 500 total count or that we removed ASVs identified as Eukaryota.

### Relative abundance of main phyla

We aggregated microbial taxa to the phylum level and divided counts by the total count per sample in order to obtain relative abundances. We then plotted the relative abundance of the main five phyla (classifying the remaining phyla as ”other”) in all fecal samples or sigmoid biopsy samples.

### Alpha-diversity

We computed the Shannon and Simpson indices with the *plot richness* function from the Phyloseq package. For datasets with two time points [24, 19], only the samples from the first time point were used. The alpha diversity was computed separately for each sample type in datasets with both fecal and biopsy samples [28, 39].

### Differential abundance analysis

We applied three recently published methods for differential abundance (DA) testing to the microbial count table of each dataset. To uncover trends at different levels of taxonomic granularity, each method was used on each dataset five times, aggregated at the phylum, class, order, family, and genus level, respectively. We used ANCOM-BC [47] and LinDA [48] for differential abundance testing of microbial composition data, as well as the scCODA model [49], a general Bayesian modeling framework for high-throughput sequencing data. All three methods take into account that the data is inherently compositional, i.e. only proportional analyses are valid [52]. To avoid biases caused by the type of sampled microbial community, we only used stool samples for this analysis, leaving out the data from [24], as well as the microbial data from the sigmoid colon in the works of [28, 39]. Furthermore, we only analyzed the samples from the initial collection in the data of [19]. For all methods and taxonomic ranks, the nominal false discovery rate was set to p < 0.2, a value commonly used in DA testing of microbial populations.

While ANCOM-BC and LinDA detect compositional changes of a feature with respect to all other features, the scCODA model requires a reference feature that is present in most samples and has low dispersion over all samples [49]. To ensure comparability, we chose the same reference on the Genus level for all datasets, and further enabled comparability across taxonomic ranks by using the reference’s ancestor at a every taxonomic rank as the reference feature for the respective aggregation level. We further did not consider genera from the Bacteroidota or Firmicutes phyla as references, as these are likely to contain many differentially abundant genera, leaving us with three candidate genera that are present in every dataset, which we selected for further inspection (Table S5). For *Escherichia/Shigella*, the dispersion was very high (> 0.3) in three datasets ([39, 28, 38]), while *Slackia* was never found in more than 20% of samples in any dataset. The remaining genus *Parasutterella* is rare in only two datasets ([13, 39]), and has rather low dispersion (< 0.055) in all datasets. Furthermore, the abundance of species from this genus was only associated with IBS in one previous analysis [72]. However, the LEfSe method [73] used in [72] was recently shown to produce false-positive results on amplicon sequencing data [74], leading us to the conclusion that *Parasutterella* is likely not affected by IBS and therefore suited as a reference for our purposes.

In the shared ASV analysis, we applied the three models ANCOM-BC, LinDA, and scCODA with the same method specifications as before, taking the ASV corresponding to *Parasutterella excrementihominis* as the reference for scCODA to provide comparability to the previous analyses. Each method was used to determine differentially abundant ASVs in the two data subsets consisting of only samples from either the Nagel [15] or Pozuelo [19] datasets, and the combined data with samples from both sources. For LinDA, we additionally ran a fourth model, in which we also adjusted for the data source in the combined data by adding random effects for the source datasets.

### Classification analysis on shared ASVs

Besides applying DA testing, we used a classification approach to determine the relationship between IBS and ASVs. For this part of the analysis, we looked at the ASVs shared between the Nagel and Pozuelo datasets. We used only the observations from the first time point in the longitudinal analysis, similar to the DA testing approach. We also excluded ASVs that occurred in less than 10% of the observations to avoid association with spurious taxa. In addition, we removed all ASVs not belonging to the Kingdom of bacteria. These preprocessing steps reduced the number of shared ASVs from 806 (see Fig. 5A) to 373 (see Fig. 5B). Of 209 observations, 30 observations were from the Nagel [15] and 179 from the Pozuelo [19] dataset. 81 subjects were diagnosed with IBS, and 128 healthy controls. To determine the relationship between the dependent variable health status and ASVs as independent variables, we used a sparse log-contrast model for classification with the tuning parameter λ from the trac package [75]. This modeling approach accounts for the compositional structure of the covariates. A log transformation is required for the model, and a pseudo-count of 1 was added to avoid log(0). The explicit model formulation is

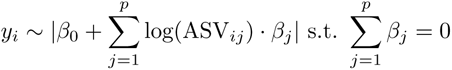

where *y_i_* 2 {Healthy (—1), IBS (1)}, *i* = 1, …, *n* indicate the subject, and ASV*_j_* denotes the count of the *j*-th ASV. We used 5-fold cross-validation to determine the optimal tuning parameter of the model, applying the heuristic of setting the tuning parameter ”1 standard error” away from the minimum misclassification error.

### High-dimensional compositional mean test

We employed the high-dimensional compositional mean test [45] to test for a shift in the compositional mean. The test builds upon the centered log-ratio transformation (CLR) [44] to overcome the constraints imposed by compositional data and tests whether the mean of the CLR transformatioed data differ across two groups. Since the CLR transformation cannot handle zeros, we replaced them with a pseudo-count of 1. The test was applied to the compositions on the Family level to examine two aspects: (1) Pairwise similarity of the mean compositions across different datasets and (2) compositional mean shift between IBS and healthy within the different datasets. For the second part, we focused on taxa that appeared in at least three datasets.

## Author’s contributions

SC researched the studies to be included, performed the data preprocessing and initial exploratory data analyses. JO conducted the differential abundance testing. VT performed the classification analysis on shared ASVs and compositional mean test. SC, JO, VT and CLM wrote the manuscript. MM and CLM provided advice and guidance throughout the project.

## Acknowledgements

We thank sincerely Dr. Robyn Nagel for sharing with us their raw data and metadata, and allowing us to share them publicly in this meta-analysis. In addition, we thank Dr.Labus, Prof.Ostrowski and Dr.Kulecka for sharing with us additional covariates on the Labus and Zeber-Lubecka datasets. We thank Nathan Fox who extracted IBS subtype metadata from the SRA website for the Pozuelo dataset.

