## Supplementary Materials for "MetaIBS - large-scale amplicon-based meta analysis of irritable bowel syndrome"

**Table S1** Demographic parameters of samples included in meta-analysis

| | **Healthy controls** | **IBS** | |
|---|---|---|---|
| **Labus et al., 2017** | | | |
| Number of individuals | 23 | 29 | |
| Age in years, mean (SD) | 26.0 (6.5) | 27.8 (11.9) | |
| Female, n (%) | 14 (61%) | 21 (72%) | |
| BMI, mean (SD) | 25.0 (4.8) | 24.0 (4.7) | |
| **Lo Presti et al., 2019** | | | |
| Number of individuals | 30 | 23 | |
| Age in years, mean (SD) | 51.2 (9.2) | 48.9 (8.7) | |
| Female, n (%) | 12 (40%) | 17 (74%) | p=0.03 |
| **AGP, 2021** | | | |
| Number of individuals | 594 | 589 | |
| Age in years, mean (SD) | 46.0 (12.8) | 45.7 (13.3) | |
| BMI, mean (SD) | 23.7 (2.8) | 23.7 (3.6) | |
| **Liu et al., 2020** | | | |
| Number of individuals | 44 | 84 | |
| Age in years, mean (SD) | 39.0 (12.9) | 42.0 (12.0) | |
| Female, n (%) | 19 (43%) | 29 (35%) | |
| BMI, mean (SD) | 23.8 (3.7) | 23.5 (3.5) | |
| **Hugerth et al., 2020** | | | |
| Number of individuals | 287 | 85 | |
| Age in years, mean (SD) | 47.6 (12.5) | 45.8 (12.2) | |
| Female, n (%) | 164 (57%) | 53 (62%) | |
| BMI, mean (SD) | 24.6 (3.9) | 23.7 (3.6) | |
| **Mars et al., 2020** | | | |
| Number of individuals | 12 | 27 | |
| Age in years, mean (SD) | 40.5 (13.3) | 38.6 (12.6) | |
| Female, n (%) | 8 (67%) | 21 (78%) | |
| BMI, mean (SD) | 26.9 (3.5) | 28.3 (8.2) | |
| **Zhu et al., 2019** | | | |
| Number of individuals | 14 | 15 | |
| Age in years, mean (SD) | 28.5 (6.2) | 47.7 (14.7) | p=0.001 |
| Female, n (%) | 6 (43%) | 5 (33%) | |
| **Nagel et al., 2016** | | | |
| Number of individuals | 15 | 15 | |
| Age in years, mean (SD) | 39.5 (13.2) | 44.7 (13.8) | |
| Female, n (%) | 10 (67%) | 12 (80%) | |
| **Zeber-Lubecka et al., 2016** | | | |
| Number of individuals | 17 | 73 | |
| Female, n (%) | *NA* | 55 (75%) | |

**Table S2** Data processing for all datasets

| Studies | Sequencing | Demultiplexing | Primers found | Primers filtering/trimming | Quality cutoff | Min length | Error rate model adjustment | ASVs discarded | Bloom sequences discarded | Samples discarded |
|---|---|---|---|---|---|---|---|---|---|---|
| Labus et al., 2017 | single-end | - | REV | Filt+Trim | 10 | 150 | - | 2 | - | 0 |
| Lo Presti et al., 2019 | single-end | - | FWD | Filt+Trim | 10 | 150 | - | 0 | - | 106 |
| Ringel-Kulka et al., 2015 | single-end | - | Both | Filt+Trim | 10 | 150 | - | 5 | - | 1 |
| AGP, 2021 | single-end | - | None | - | 10 | 150 | - | 1,576 | 55 | 43 |
| Liu et al., 2020 | single-end | - | REV | Filt+Trim | 10 | 150 | - | 16 | - | 0 |
| Pozuelo et al., 2015 | single-end | - | REV | Filt+Trim | 10 | 150 | - | 477 | - | 0 |
| Fukui et al., 2020 | paired-end | - | Both | Filt+Trim | 10 | 150 | - | 4 | - | 1 |
| Hugerth et al., 2020 | paired-end | - | Both | Filt+Trim | 10 | 150 | - | 16 | - | 79 |
| Mars et al., 2020 | paired-end | - | Both | Filt+Trim | 10 | 150 | - | 3 | - | 3 |
| Zhu et al., 2019 | paired-end | - | FWD | - | 10 | 150 | - | 0 | - | 0 |
| Zhuang et al., 2018 | paired-end | - | Both | Filt+Trim | 10 | 150 | - | 0 | - | 0 |
| Nagel et al., 2016 | single-end | Yes | Both | Filt+Trim | 10 | 150 | Yes | 3 | - | 0 |
| Zeber-Lubecka et al., 2016 | single-end | - | Primer mix | Trim first 15bp | 10 | 150 | Yes | 10 | - | 0 |

Demultiplexing indicates whether samples had to be demultiplexed.

Primers found column indicates whether primers were found in the reads and removed.

Primers filtering/trimming indicates whether only samples containing the primers were kept (Filt) and/or if the primer(s) found were trimmed off (Trim). For the Zhu dataset, primers were in the middle of the reads, and thus could not be trimmed off.

Quality cutoff column indicates the quality score at which reads were truncated.

Min length column indicates the length (bp) below which reads were discarded.

Error rate model adjustment column indicates whether the error rate model had to be manually adjusted.

ASV discarded column indicates the number of ASVs discarded because they belonged to Eukaryota or to an unknown phylum.

Bloom sequences discarded indicates the number of ASVs discarded because they were identified as bloom sequences (AGP data).

Samples discarded column indicates the number of samples discarded because they had a total count below 500.

**Table S3** Data availability

| Studies | SRA/ENA accession number | Metadata accession | Metadata available |
|---|---|---|---|
| Labus et al., 2017 | PRJNA373876 | SRA (IBS subtype shared after inquiry to author) | age, gender, BMI, IBS subtype |
| Lo Presti et al., 2019 | PRJNA391149 | SRA | age, gender, IBS subtype, sample type |
| Ringel-Kulka et al., 2015 | SRP066323 | - | - (no HC/IBS label) |
| AGP, 2021 | PRJEB11419 | SRA | age, BMI, bowel movement quality/frequency, diet, comorbidities |
| Liu et al., 2020 | PRJNA544721 | SRA | age, gender, BMI, IBS subtype (all IBS-D), Bristol stool scale, IBS symptom severity scale |
| Pozuelo et al., 2015 | PRJNA268708 | SRA | IBS subtype, collection time point (0, 1month) |
| Fukui et al., 2020 | PRJNA637763 | SRA | - |
| Hugerth et al., 2020 | PRJEB31817 | SRA | age, gender, BMI, psychological distress |
| Mars et al., 2020 | PRJEB37924 | SRA | age, gender, BMI, IBS subtype, collection time point (0, 6months) |
| Zhu et al., 2019 | PRJNA566284 | SRA | age, gender |
| Zhuang et al., 2018 | SRP150089 | SRA | IBS subtype (all IBS-D) |
| Nagel et al., 2016 | Private inquiry to author | Private inquiry to author | age, gender, IBS subtype (all IBS-D) |
| Zeber-Lubecka et al., 2016 | PRJEB11252 | SRA (gender shared after inquiry to author) | gender, IBS subtype |

**Table S4** Number of samples before and after pre-processing of the data.

| **Studies** | **Number of individuals recruited (HC/IBS)** | **Number of samples downloaded (HC/IBS)** | **Number of samples analyzed (HC/IBS)** | **Notes** |
|---|---|---|---|---|
| Labus et al., 2017 | 23/29 | 23/29 | 23/29 | - |
| Lo Presti et al., 2019 | 47/44 | stool:40/36<br>sigmoid:46/41 | stool:27/19<br>sigmoid:7/4 | - |
| Ringel-Kulka et al., 2015 | 20/56 | 76 | 75 | HC/IBS label unavailable |
| AGP, 2021 | 4722/645 | 645/645 | 594/589 | 645 HC samples randomly selected among the 4,722 available |
| Liu et al., 2020 | 44/84 IBS-D | 44/84 | 44/84 | - |
| Pozuelo et al., 2015 | 66/113 | t0:66/113<br>t0+1mo:22/72 | t0:66/113<br>t0+1mo:22/72 | 94 individuals provided a second fecal sample 1 month later |
| Fukui et al., 2020 | 26/85 | 26/85 | 26/84 | - |
| Hugerth et al., 2020 | 317/66 | stool:161/57<br>sigmoid:306/83 | stool:130/44<br>sigmoid:274/77 | inconsistency between number of samples reported in paper and samples available on SRA |
| Mars et al., 2020 | 14/28 | t0:12/27<br>t0+6mo:12/21 | t0:12/25<br>t0+6mo:12/20 | 33 individuals provided a second sigmoid biopsy sample 6 months later |
| Zhu et al., 2019 | 15/15 | 14/15 | 14/15 | - |
| Zhuang et al., 2018 | 13/27 IBS-D | 10/20 | 10/20 | only samples pre-rifaximin treatment were downloaded |
| Nagel et al., 2016 | 15/15 IBS-D | 15/15 | 15/15 | only Blastocystis-negative samples downloaded |
| Zeber-Lubecka et al., 2016 | 30/72 | 17/73 | 17/73 | only samples pre-rifaximin treatment were downloaded |

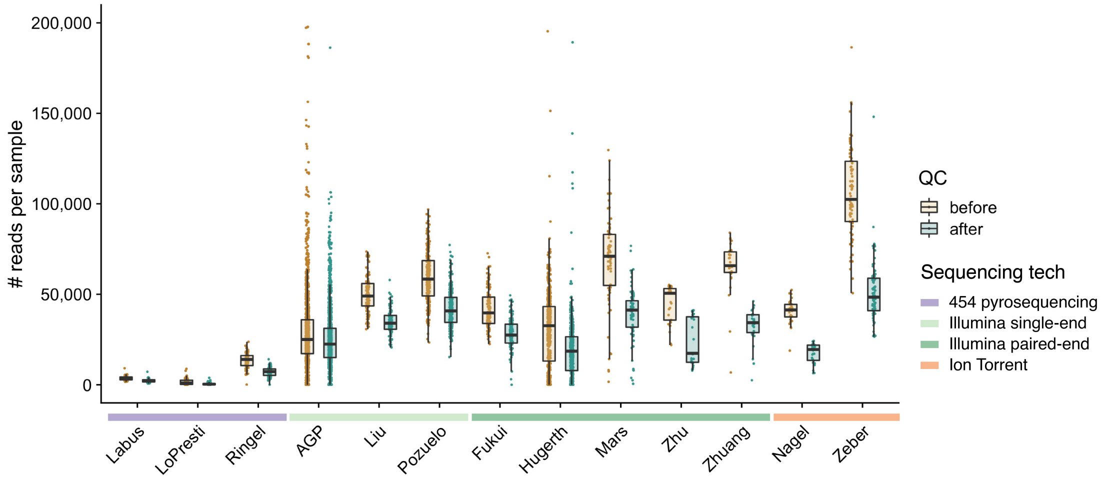


**Figure S1 Sequencing depth before and after quality control of the data.** Raw 16S rRNA sequencing data in FASTQ files from 13 datasets was quality-filtered and amplicon sequence variants (ASVs) were inferred from the reads with the DADA2 package [65]. A standardized pipeline was used to process all samples (Supplementary Table S2). This plot shows the samples' sequencing depth in the raw data (before QC) and after processing the data (after QC). Colored bars on the x axis show the sequencing technology used for each dataset.

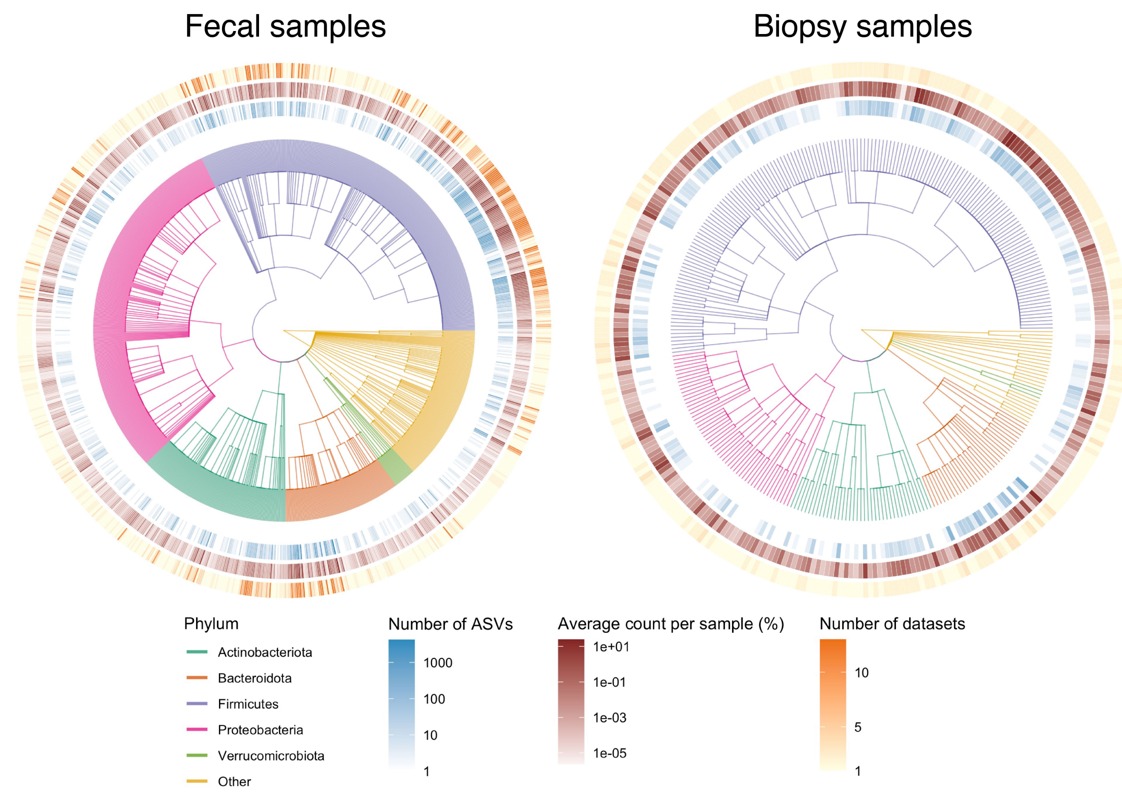


**Figure S2 Taxonomic tree of genera detected** in all fecal samples (left) and in all sigmoid mucosa biopsy samples (right). Each leaf is a genus, branches are colored by phylum. Outer rings represent the number of ASVs belonging to each genus (blue gradient), average count of each genus in a sample (brown gradient), and the number of datasets in which each genus was found (orange gradient).

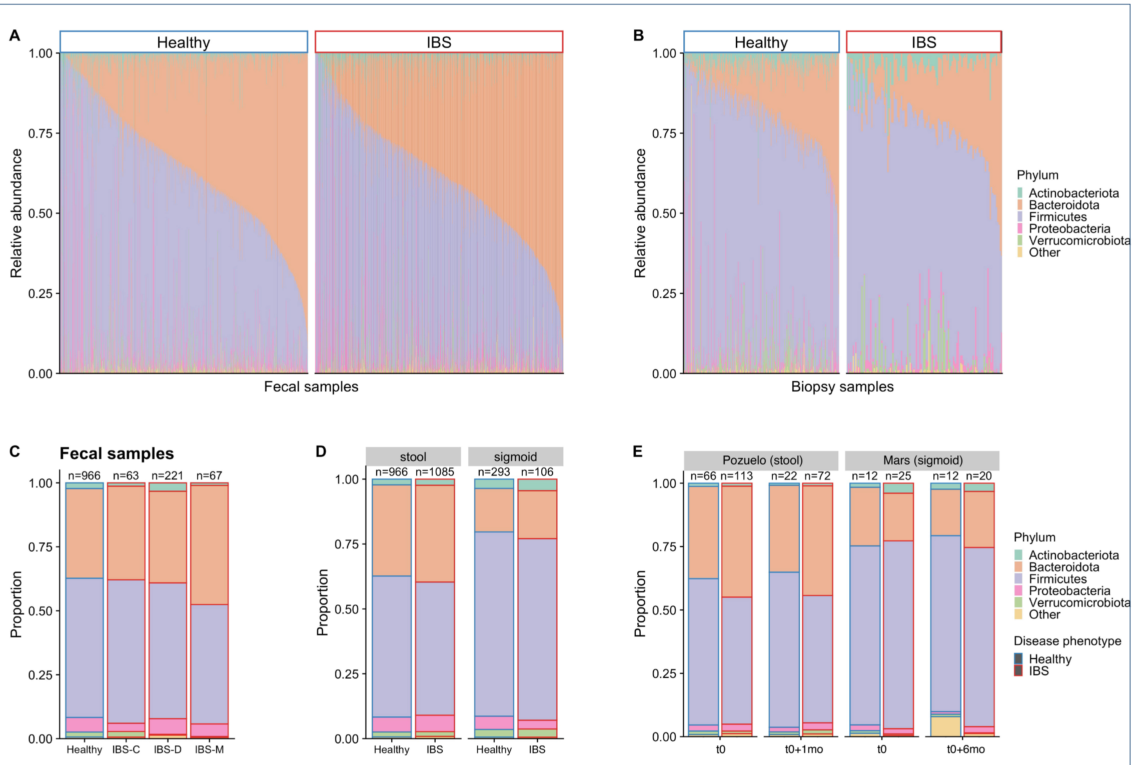


**Figure S3 Relative abundance of main phyla.** Relative abundance of main phyla in all (A) fecal samples or (B) sigmoid colon biopsy samples. (A-B) Healthy samples on the left, IBS samples on the right of each panel. Relative abundance of main phyla averaged across (C) fecal samples (only 1st collection time point for Pozuelo dataset) classified by IBS subtype (734 IBS samples of unknown subtypes were not shown); (D) all samples (only 1st collection time point for Pozuelo and Mars datasets) classified by sample type (stool, sigmoid colon) and disease phenotype (healthy, IBS); (E) samples from the Pozuelo and Mars datasets, classified by collection time point (t0, t0+1month, t0+6months) and disease phenotype (healthy, IBS). (C-E) Number of samples per category is shown above each bar.

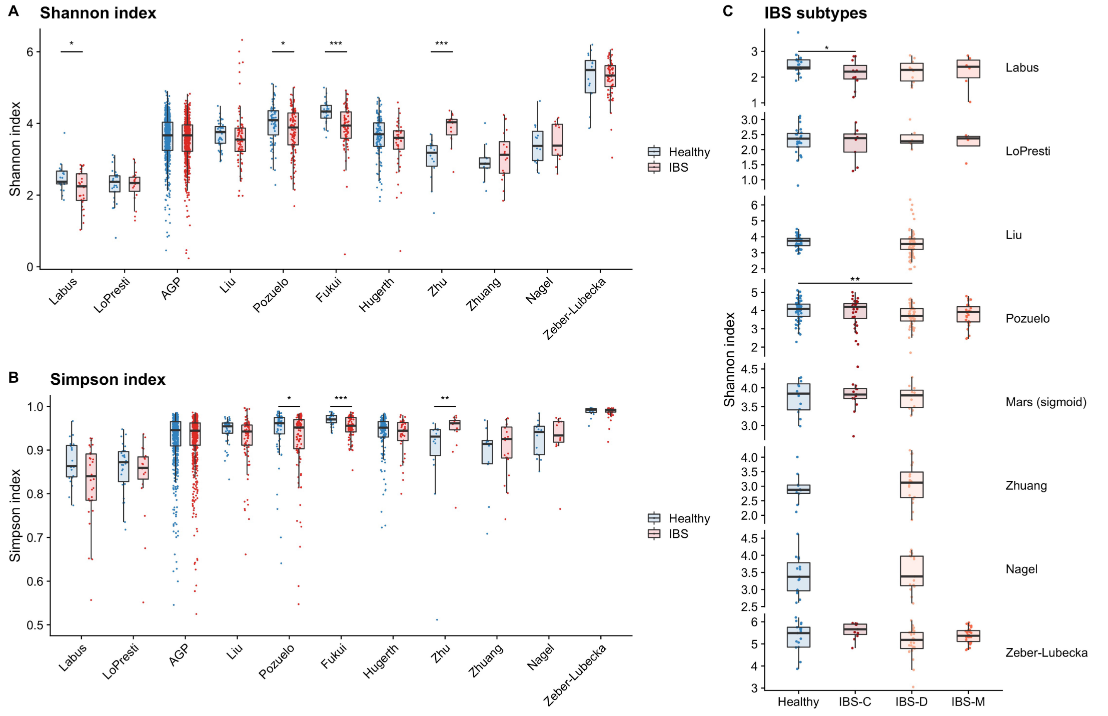


**Figure S4 $\alpha$-diversity index in healthy and IBS samples across datasets.** Left panels show the (A) Shannon and (B) Simpson indexes in stool samples across datasets (for Pozuelo, only the first collection time point is shown). (C) Shannon index in datasets with information on IBS subtype (for Pozuelo and Mars, only the first collection time point is shown; for Lo Presti only stool samples are shown). Statistical significance determined by a two-way wilcoxon test (* p<0.05 ; ** p<0.01; *** p<0.001).

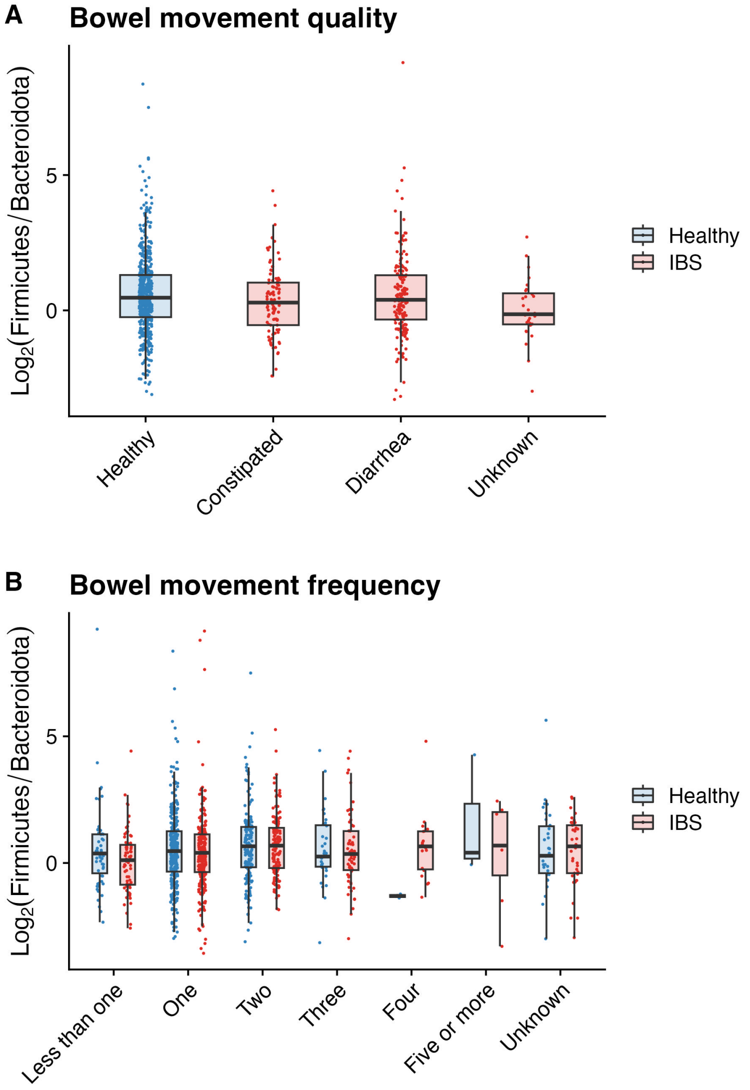


**Figure S5 Log ratio of Firmicutes:Bacteroidota in the AGP dataset**. (A) Healthy samples with normal bowel movement quality and IBS samples with either constipation, diarrhea or unknown bowel movement quality show similar F/B ratio. (B) F/B ratio in healthy and IBS samples based on the reported number of daily bowel movements reported.

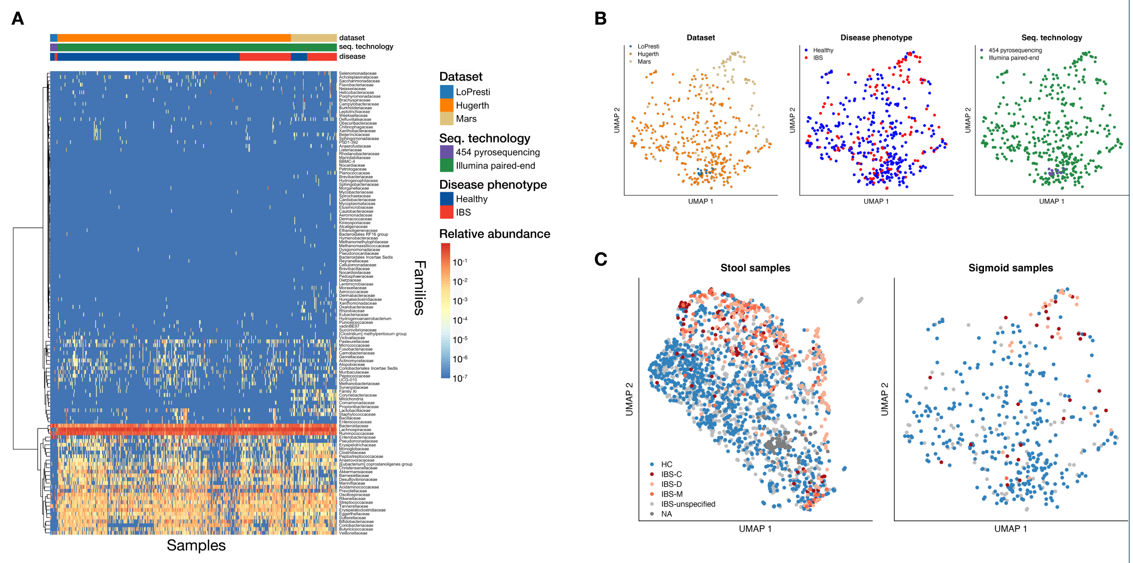


**Figure S6 Microbial samples from biopsies of the sigmoid colon do not cluster by disease phenotype**. (A) Heatmap of microbial families abundance in all sigmoid mucosal biopsy samples. Families are in rows, clustered with the ward algorithm; samples are in columns, ordered by dataset and disease phenotype. Counts were normalized (sum per sample is 1). (B) UMAP was run on log-ratios between microbial families in all sigmoid mucosal biopsy samples. Samples are colored by dataset (left), disease phenotype (middle), or sequencing technology (right). (C) UMAP of fecal samples (left) and sigmoid mucosal biopsy samples (right), colored by IBS subtype. Samples for which the IBS subtype is unknown are colored in grey. (A-C) All sigmoid mucosal biopsy samples were plotted (n=431), including both collection time points in the Mars dataset.

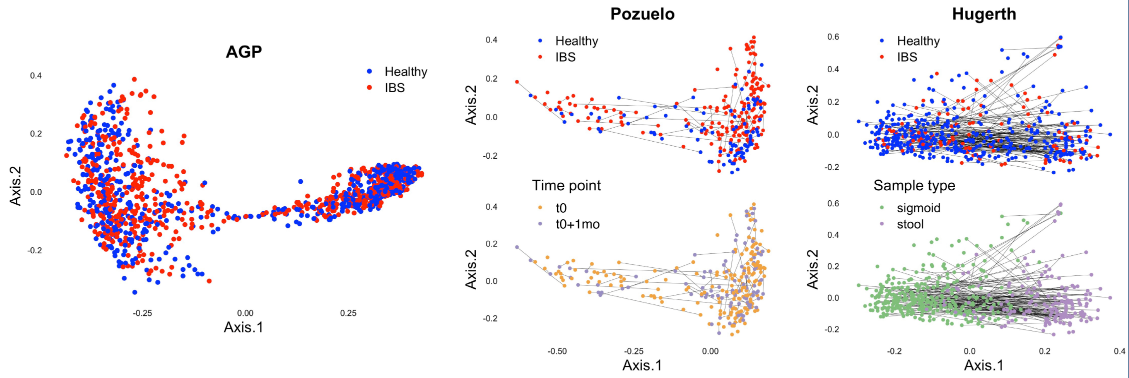


**Figure S7 In datasets with higher cohort size, samples do not cluster by disease phenotype**. Principal Coordinate Analysis (PCoA) on the AGP (left; n=1,183), Pozuelo (middle; n=273) and Hugerth (right; n=525) datasets, using the Bray-Curtis dissimilarity. Samples are colored by disease phenotype. In the Pozuelo dataset, samples are also colored by collection time point (middle bottom). In the Hugerth dataset, samples are also colored by sample type (right bottom). In the Pozuelo and Hugerth datasets, lines connect paired samples (same individual with distinct collection time points, or different sample types).

| Genus | First Author | Dispersion | Presence |
|---|---|---|---|
| *Slackia* | AGP | 0.001 | 0.143 |
| *Slackia* | Liu | 0.009 | 0.086 |
| *Slackia* | Fukui | 0.005 | 0.018 |
| *Slackia* | Zhuang | 0.001 | 0.033 |
| *Slackia* | Hugerth | 0.003 | 0.161 |
| *Slackia* | Zhu | 0.000 | 0.138 |
| *Slackia* | Labus | 0.004 | 0.019 |
| *Slackia* | Lo Presti | 0.007 | 0.022 |
| *Slackia* | Zeber | 0.002 | 0.200 |
| *Slackia* | Pozuelo | 0.000 | 0.095 |
| *Slackia* | Nagel | 0.004 | 0.200 |
| *Parasutterella* | Liu | 0.043 | 0.586 |
| *Parasutterella* | Zhuang | 0.054 | 0.500 |
| *Parasutterella* | Zhu | 0.017 | 0.207 |
| *Parasutterella* | Zeber | 0.040 | 0.667 |
| *Parasutterella* | AGP | 0.016 | 0.574 |
| *Parasutterella* | Fukui | 0.032 | 0.445 |
| *Parasutterella* | Nagel | 0.028 | 0.733 |
| *Parasutterella* | Lo Presti | 0.001 | 0.022 |
| *Parasutterella* | Labus | 0.048 | 0.096 |
| *Parasutterella* | Hugerth | 0.033 | 0.443 |
| *Parasutterella* | Pozuelo | 0.011 | 0.743 |
| *Escherichia/Shigella* | Zeber | 0.062 | 0.678 |
| *Escherichia/Shigella* | Nagel | 0.080 | 0.533 |
| *Escherichia/Shigella* | Lo Presti | 0.368 | 0.370 |
| *Escherichia/Shigella* | Zhu | 0.012 | 0.414 |
| *Escherichia/Shigella* | Labus | 0.009 | 0.038 |
| *Escherichia/Shigella* | Zhuang | 0.038 | 0.833 |
| *Escherichia/Shigella* | Liu | 0.165 | 0.938 |
| *Escherichia/Shigella* | Hugerth | 0.331 | 0.552 |
| *Escherichia/Shigella* | Fukui | 0.025 | 0.209 |
| *Escherichia/Shigella* | AGP | 0.310 | 0.117 |
| *Escherichia/Shigella* | Pozuelo | 0.190 | 0.682 |

**Table S5** Candidate genera for reference selection in scCODA that are present in all stool datasets and are not from the Bacteroidota or Firmicutes phyla. Dispersion and Presence were calculated over all stool samples (only initial collection for [19]) in the respective datasets.

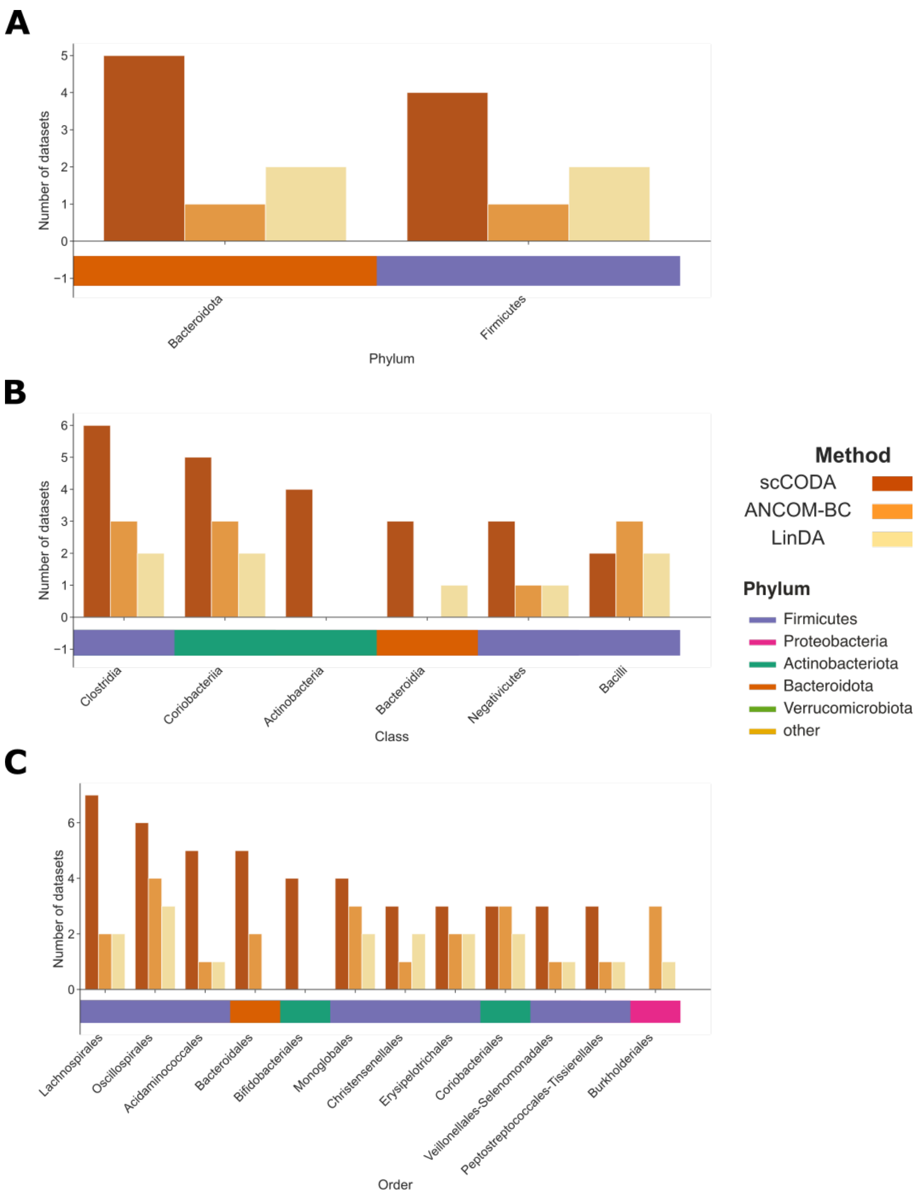


**Figure S8 Differential abundance testing of stool samples from 11 different datasets**, aggregated on the (A) phylum, (B) class, and (C) order level. Bars indicate the number of datasets where a taxonomic group was found to be differentially abundant. Only genera that were differentially abundant for at least three datasets are shown. Colored boxes below the x-axis show the corresponding phylum for each group.

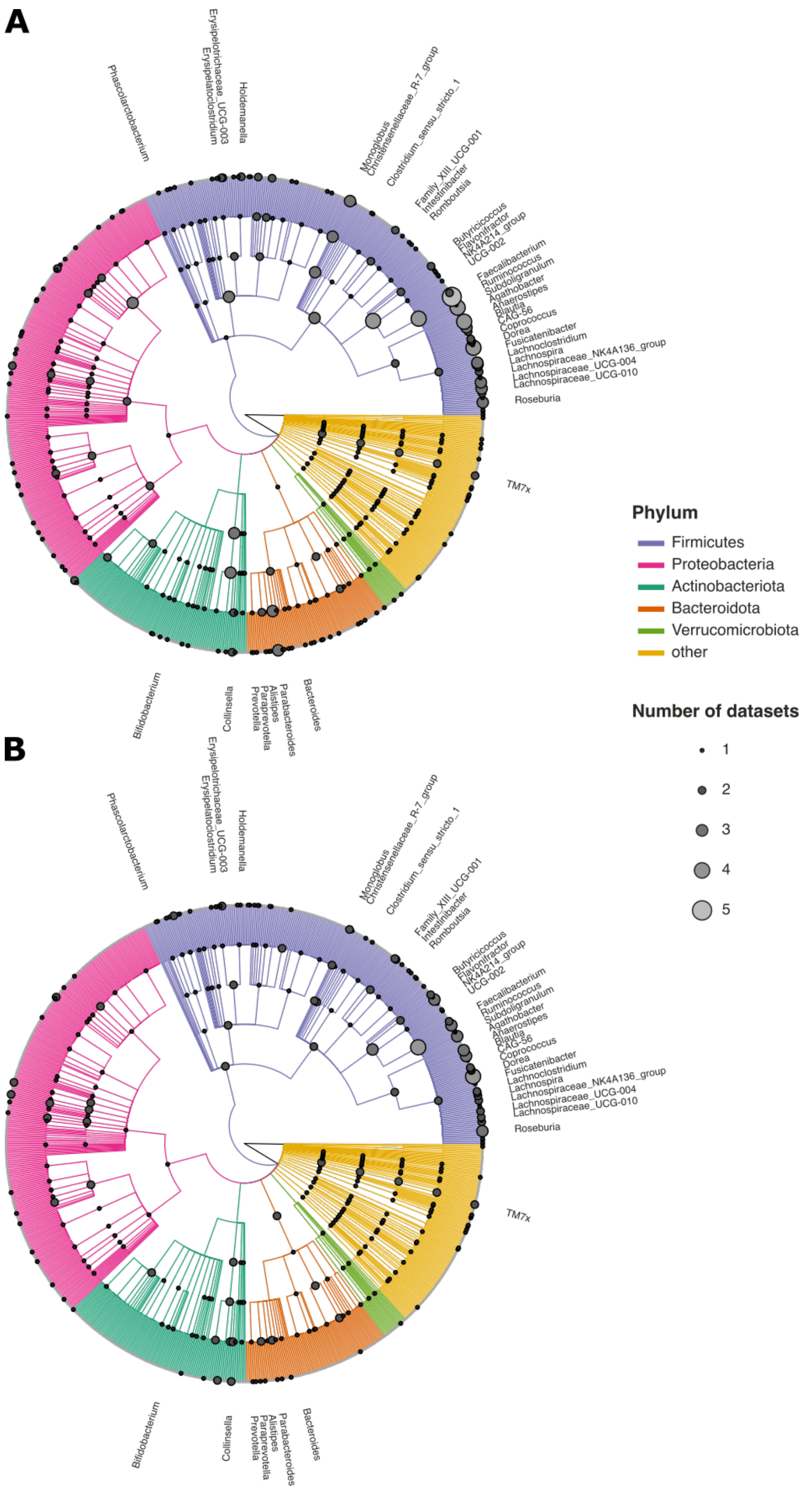


**Figure S9 Differential abundance testing visualized as taxonomic trees** including all genera that are present in at least one dataset (stool samples only). Grey circles on the nodes and leaves indicate the number of datasets that a taxon was differentially abundant in, as determined by (A) ANCOM-BC and (B) LinDA. The branch colors indicate the phylum, tip labels denote the 38 genera found to be differentially abundant in at least three datasets.
